# Persister cells and their effects on cancer stabilization therapy

**DOI:** 10.64898/2026.10.15.759390

**Authors:** Mohammadreza Satouri, Philipp M. Altrock, Jafar Rezaei, Kateřina Staňková

**Affiliations:** Institute for Health Systems Science, Faculty of Technology, Policy and Management, Delft University of Technology, Delft, The Netherlands; Department of Hematology and Oncology, University Hospital Schleswig-Holstein, Kiel University, Kiel, Germany; Department of Engineering, Systems and Services, Faculty of Technology, Policy and Management, Delft University of Technology, Delft, The Netherlands

**Keywords:** Persister cancer cells, Evolutionary game theory, Stability analysis, Bifurcation analysis, Drug resistance

## Abstract

Some cancers have the ability to persist under treatment. Persister cancer cells can be a small subpopulation of cancer cells that enter a dormant or quiescent state, thereby evading the effects of chemotherapy, targeted therapy, or immunotherapy. Motivated by these observations, we develop and investigate an evolutionary game theory model that captures the interactions between three types of cancer cells: treatment-sensitive, drug-tolerant persister, and treatment-resistant cells. In our model, persister cells transition into resistant cells at a rate that can depend on the treatment intensity. We analyze the model’s equilibria and derive conditions for their existence and stability. In particular, we show that an equilibrium in which sensitive and resistant cells coexist is stable when intra-type competition is stronger than inter-type competition. We also show that equilibria in which only one cell type persists require the transition rate to exceed an explicit threshold determined by the treatment dose and competition parameters. Our findings reveal that persister cells can both stabilize and destabilize tumor dynamics, depending on treatment and cell type-transition rates, highlighting the importance of accounting for persister behavior in designing effective evolutionary therapies.

## 1 Introduction

Cancer resistance to therapy is a multifaceted challenge, encompassing both intrinsic and acquired mechanisms that enable tumor cells to evade treatment [2, 32, 15]. In chemotherapy, resistance can arise through a variety of mechanisms, including, but not limited to, increased drug efflux, enhanced DNA repair, evasion of apoptosis, and epigenetic silencing of tumor suppressor genes [21, 16, 14, 38]. In addition, the tumor microenvironment can create protective niches that shield cancer cells from therapeutic agents, further complicating treatment outcomes [31, 34]. Resistance to targeted therapies and immunotherapy presents additional challenges due to the specificity of drug targets and the adaptability of tumor signaling pathways [1]. Such resistance can arise through mutations in drug targets or activation of alternative signaling pathways [30]. A particularly important contributor to relapse and treatment failure is the pre-treatment existence of drug-tolerant persister cell (DTP) populations. This cell population can survive initial therapy and may later acquire permanent resistance [6, 28].

DTPs represent a small subpopulation of cancer cells that survive therapeutic interventions by entering a transient, non-proliferative state [35]. Unlike cells that acquire resistance through genetic mutations, DTPs evade treatment via reversible, non-genetic adaptations, such as epigenetic reprogramming [17], metabolic shifts [23], and phenotypic plasticity [18, 25]. These cells have been identified in various cancers, including melanoma, breast, lung, and pancreatic cancers [6, 11, 23, 13]. Their ability to survive initial treatments and later resume proliferation contributes significantly to cancer relapse and poses a major challenge in oncology [6]. DTPs tolerate treatment through various mechanisms, including stem cell-like properties, such as self-renewal and plasticity [3].

Understanding and targeting the unique biology of DTPs is crucial for developing more effective cancer treatments and improving patient outcomes [25, 5]. One promising direction is to incorporate persister dynamics into evolutionary therapies, an emerging paradigm that leverages Darwinian evolutionary principles to manage cancer progression and treatment resistance [9, 10, 8]. Unlike traditional approaches that aim for maximum tumor eradication, evolutionary therapy seeks to maintain a stable tumor burden by preserving a population of treatment-sensitive cells to suppress the growth of resistant ones through competitive interactions [8, 36]. This strategy, also known as adaptive therapy, involves modulating drug dosing and scheduling to exploit the fitness costs associated with resistance [8, 39]. However, a critical gap remains: while prior studies have examined persister cell dynamics through empirical observation or standard mathematical models [6, 28], these frameworks rarely integrate DTPs into adaptive or evolutionary therapy designs. Because DTPs survive initial therapy and serve as a reservoir for the eventual emergence of fully resistant clones [12], they add a critical dimension to treatment dynamics. In this work, we address this gap by extending evolutionary therapy approaches to explicitly include persister dynamics.

We develop and investigate a competitive Lotka-Volterra mathematical model that includes competition among sensitive, persister, and resistant cells, and a transition or mutation rate from persister to resistant cells. We analyze how persister cells alter long-term dynamics under constant treatment. In the remainder of the manuscript, we use the term mutation rate to refer to the rate of cells switching from the persister to a resistant state.

The remainder of this paper is organized as follows. Section 2 introduces the mathematical model governing the three-dimensional tumor dynamics. Section 3 provides a comprehensive stability analysis of all possible biological equilibria. Section 4 illustrates typical cancer trajectories and the distribution of these equilibria under varying mutation rates and treatment doses. Finally, Section 5 discusses the implications of our findings for evolutionary therapy design.

## 2 Mathematical Model

To capture the complex evolutionary dynamics of cancer cell populations using a therapeutic agent (*m*), we employ a three-dimensional dynamical system based on the competitive Lotka-Volterra equations [37, 19]. This framework models interactions among three distinct subpopulations of tumor cells: drug-sensitive cells (with population *x*_*S*_), drug-resistant cells (with population *x*_*R*_), and drug-tolerant persister cells (with population *x*_*P*_). Let ***x*** be the vector including the 3 populations, i.e., ***x*** = (*x*_*S*_, *x*_*R*_, *x*_*P*_)^*T*^. Each subpopulation is governed by a differential equation that captures its growth, apoptosis, and competition within and between cell types, represented by *α*_*ij*_ with *i, j ∈ {S, R, P}* for space and resources. The competition matrix, which consists of the competition coefficients between different cell types, is expressed as

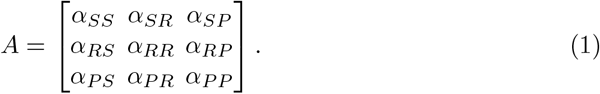

Moreover, the model incorporates a possibility for persister cells to mutate to resistant ones, representing the ability of persisters to acquire stable, heritable resistance over time through genetic, epigenetic, and phenotypic changes. We assume the following evolution of the population of three types of cancer cells:

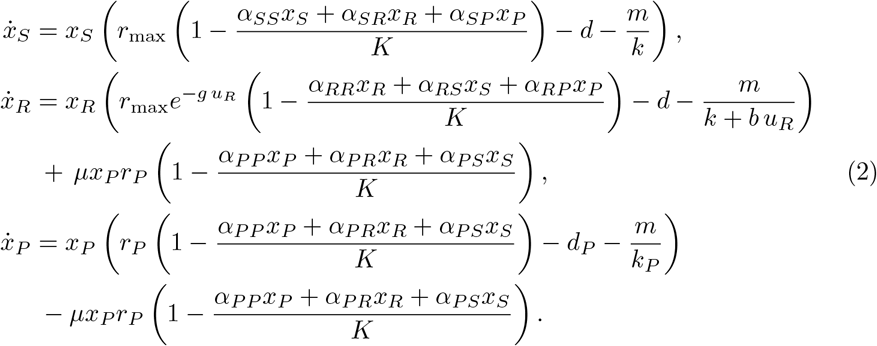

In (2), it is assumed that cancer cells can be eliminated either by natural cell death *d* and *d*_*P*_, or treatment. The efficacy of the treatment is reduced by pre-existing drug resistance *k*, and the benefit *b* of the resistance *u*_*R*_.

The growth rates for sensitive and persister cells are *r*_max_ and *r*_*P*_, respectively. We assume that resistance has a cost expressed through growth rate 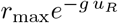 of resistant cells decreasing with resistance, and *g* is the magnitude of the cost of resistance.

Persister cells mutate to resistant ones at the mutation rate *µ*, which can be a function of treatment or a stochastic variable. Here, we assume that *µ∈* (0, 1).

Table 1 summarizes the parameters in (2). Numerical values used in the simulations are given separately in Table 2 and in the corresponding figure captions.

**Table 1:** Parameters of the cancer population dynamics in (2).

| Parameter | Description | Range |
| --- | --- | --- |
| $r_{\max}$ | Maximum growth rate of sensitive and resistant cells. | $\mathbb{R}_0^+$ |
| $r_P$ | Growth rate of persister cells. | $\mathbb{R}_0^+$ |
| $K$ | Carrying-capacity. | $\mathbb{R}_0^+$ |
| $\alpha_{ij}$ | Competition coefficient. | $\mathbb{R}_0^+$ |
| $d$ | Natural death rate of sensitive and resistant cells. | $\mathbb{R}_0^+$ |
| $d_P$ | Natural death rate of persister cells. | $\mathbb{R}_0^+$ |
| $m$ | Treatment dose. | $\mathbb{R}_0^+$ |
| $k$ | Innate resistance of sensitive and resistant cells. | $[1, +\infty]$ |
| $k_P$ | Innate resistance of persister cells. | $[1, +\infty]$ |
| $u_R$ | Resistance rate of resistant cells. | $\mathbb{R}_0^+$ |
| $b$ | Benefit of resistance under treatment. | $\mathbb{R}_0^+$ |
| $g$ | Cost of resistance. | $\mathbb{R}_0^+$ |
| $\mu$ | Mutation rate from persister to resistant cells. | $(0, 1)$ |

**Table 2:**
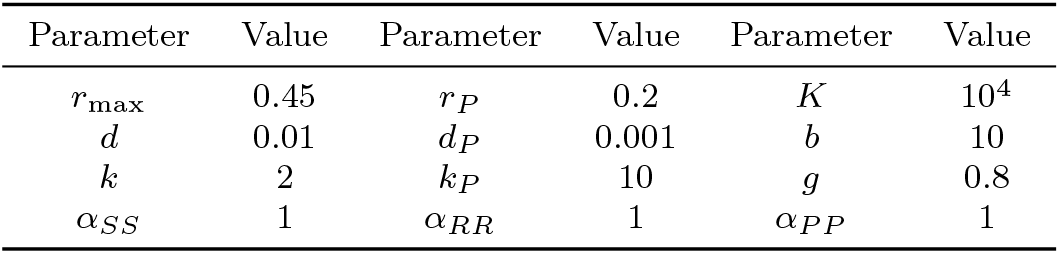
Parameters of cancer dynamics (2) utilized in our simulations.

| Parameter | Value | Parameter | Value | Parameter | Value |
| --- | --- | --- | --- | --- | --- |
| $r_{\max}$ | 0.45 | $r_P$ | 0.2 | $K$ | $10^4$ |
| $d$ | 0.01 | $d_P$ | 0.001 | $b$ | 10 |
| $k$ | 2 | $k_P$ | 10 | $g$ | 0.8 |
| $\alpha_{SS}$ | 1 | $\alpha_{RR}$ | 1 | $\alpha_{PP}$ | 1 |

## 3 Stability of different equilibria

We analyze the stability of all equilibria of dynamics (2) using the Lyapunov indirect method [20]. The Jacobian matrix of the system

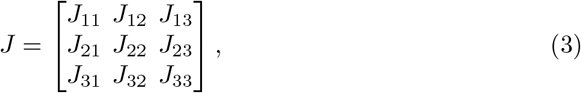

contains elements

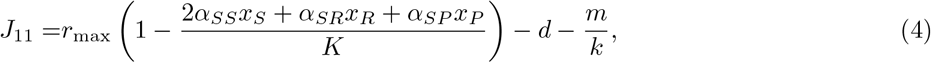

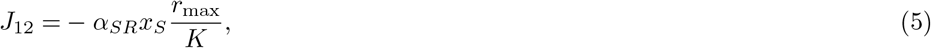

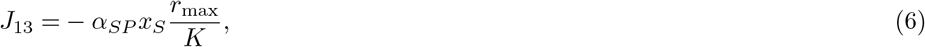

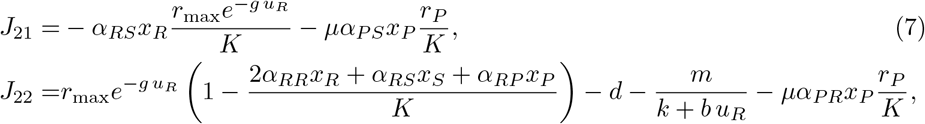

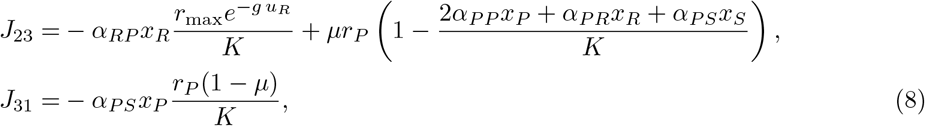

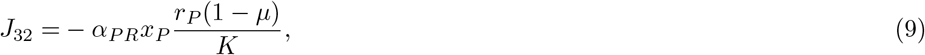

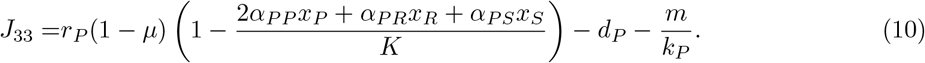

### 3.1 Trivial Equilibrium

Let ***x***^⋄^ denote a trivial equilibrium, i.e., ***x***^⋄^ = (0, 0, 0)^⊤^.

Eigenvalues of the Jacobian (3) evaluated at *x*^⋄^ are:

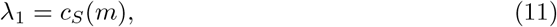

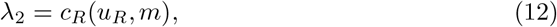

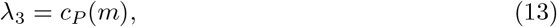

where

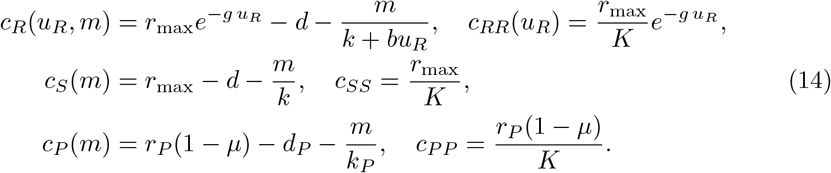

#### Remark 1.

*Note that c*_*P*_ (*m*) *also depends on µ, which is treated as a fixed parameter throughout the stability analysis*.

Through the eigenvalues (11)-(13), we can express the following theorem.

#### Theorem 1.

*The trivial equilibrium* ***x***^⋄^ *always exists. If c*_*S*_(*m*) *<* 0, *c*_*R*_(*u*_*R*_, *m*) *<* 0, *and c*_*P*_ (*m*) *<* 0, *it is locally asymptotically stable*.

*Proof*. The trivial equilibrium ***x***^⋄^ is locally stable if the eigenvalues (11)-(13) have negative real parts, implying also that *c*_*S*_(*m*) *<* 0, 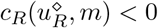, and *c*_*P*_ (*m*) *<* 0. □

#### 3.1.1 Bifurcation analysis

The eigenvalues at the TE are real. Hence, a Hopf bifurcation cannot occur at this equilibrium. A simple zero eigenvalue is possible on

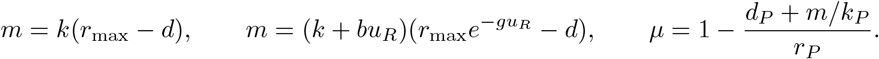

Only values in the parameter domain are relevant. The first two boundaries correspond to collisions with the FSE and FRE, respectively. Their logistic quadratic coefficients are negative, so these are transcritical bifurcations when the remaining eigenvalues are nonzero. The third boundary can connect the TE to a CRP branch. If *c*_*R*_ (*u*_*R*_, *m*) *<* 0, the linear resistant response to a small persister population is *µr*_*P*_ */*(*−c*_*R*_(*u*_*R*_, *m*)) *>* 0, and the quadratic coefficient in the persister equation is

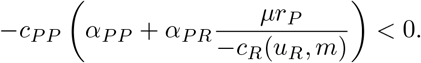

Thus this collision is transcritical when *c*_*S*_, *c*_*R*_(*u*_*R*_, *m*) ≠0, *c*_*R*_(*u*_*R*_, *m*) *<* 0, and the crossing lies in 0 *< µ <* 1. The stationary equations may exhibit a formal equilibrium bifurcation involving a branch with a negative population. That portion of the branch lies outside the biologically admissible region and cannot represent a transition between biologically feasible equilibria. Simultaneous zero eigenvalues require a separate analysis with more than one varying parameter.

In the sequel, to study whether a small persister population can grow near an equilibrium with *x*_*P*_ = 0, we calculate the coefficient of the quadratic term in the equation for *x*_*P*_. We perform this calculation at the critical mutation rate, where the persister eigenvalue is zero and the other eigenvalues are nonzero. Near this equilibrium, changes in the sensitive and resistant populations can be expressed in terms of *x*_*P*_. These local relationships, called the center manifold, describe the dynamics associated with the zero eigenvalue.

The derivatives *dx*_*S*_*/dx*_*P*_ and *dx*_*R*_*/dx*_*P*_ describe how the sensitive and resistant populations change with *x*_*P*_ along these relationships. They are obtained by solving

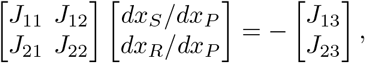

where all Jacobian entries and derivatives are evaluated at the critical equilibrium. Substituting these derivatives into the persister equation gives the following coefficient of 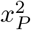:

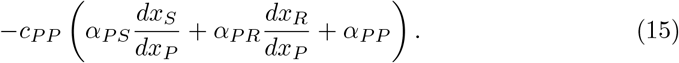

This coefficient describes the leading nonlinear effect on persister growth at the critical point. If it is nonzero and the persister eigenvalue changes with mutation at a nonzero rate, the conditions for a nondegenerate transcritical bifurcation are satisfied. If the quadratic coefficient is zero, further analysis of higher-order terms is required before the bifurcation can be classified.

### 3.2 Fully Sensitive Equilibrium

At this equilibrium, which we denote by 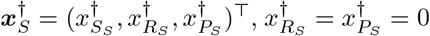, while 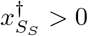. According to (2), from 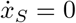 one can obtain:

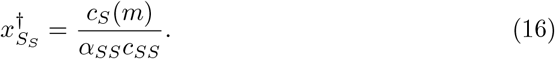

Since 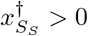, it implies *c*_*S*_(*m*) *>* 0.

By substituting 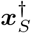 in the Jacobian matrix (3), we obtain the following matrix.

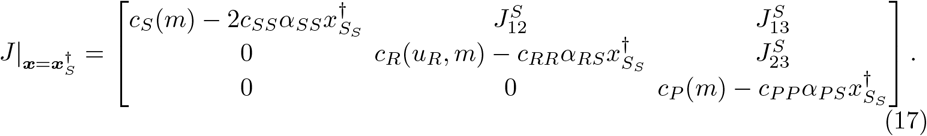

Since (17) is an upper triangular matrix, its eigenvalues are the diagonal elements of (17). The 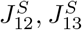, and 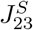 are the values of *J*_12_, *J*_13_, and *J*_23_ entries of the Jacobian (3) calculated at 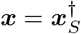. These values have no effect on the eigenvalues. By substituting the value of 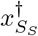 in (17), the eigenvalues become

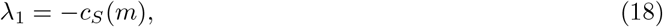

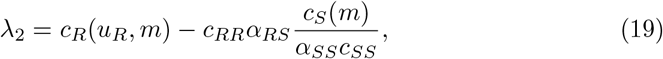

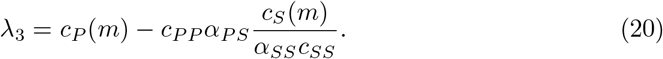

#### Theorem 2.

*If* 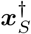 *exists and, additionally*, 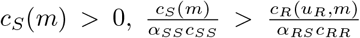, *and* 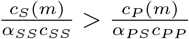, *then* 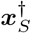*is locally asymptotically stable*.

*Proof*. According to (18), if *c*_*S*_(*m*) *>* 0, then *λ*_1_ *<* 0.

By dividing (19) and (20) by *α*_*RS*_*c*_*RR*_ and *α*_*PS*_*c*_*PP*_ respectively, we can see that *λ*_2_ and *λ*_3_ are negative when 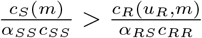 and 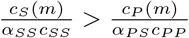. □

#### 3.2.1 Bifurcation analysis

The FSE exists for *c*_*S*_(*m*) *>* 0 and has only real eigenvalues. It cannot undergo a Hopf bifurcation. The boundary *c*_*S*_(*m*) = 0 connects it to the TE. The resistant invasion boundary is

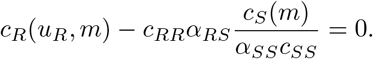

It is independent of *µ* and connects the FSE to the CSR branch when the remaining eigenvalues are nonzero and 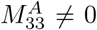. Varying mutation alone cannot cross this boundary. The persister invasion boundary is *λ*_3_ = 0 in (20). Its derivative is

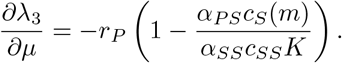

At an interior crossing with *d*_*P*_ + *m/k*_*P*_ *>* 0, this derivative is negative. Equation (15) checks the second nondegeneracy condition. A nonzero quadratic coefficient gives a transcritical collision with CAT, provided that the adjacent populations are positive.

### 3.3 Fully resistant equilibrium

At this equilibrium, which we denote by 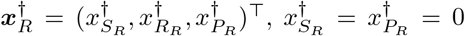 while 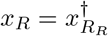 and from 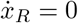:

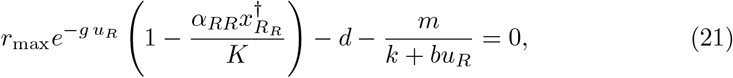

and consequently

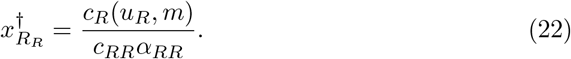

Since 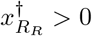, it is equivalent to *c*_*R*_(*u*_*R*_, *m*) *>* 0.

By substituting 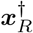 into the Jacobian matrix (3), we obtain the following matrix.

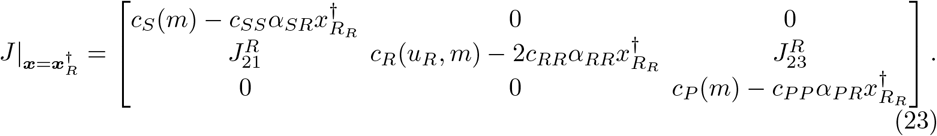

Using Laplace expansion on the first row of (23) for calculating the determinant of 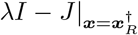, we can see that the eigenvalues are the diagonal elements of (23), and 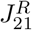, and 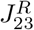 have no effect on the eigenvalues. By substituting the value of 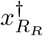 in (23), the eigenvalues become

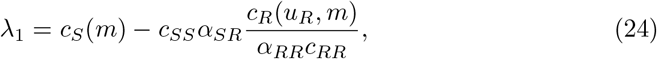

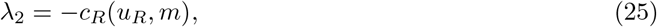

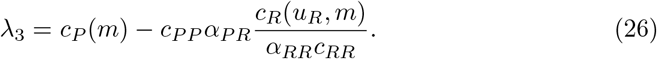

#### Theorem 3.

*If* 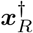 *exists, and additionally* 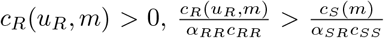, and 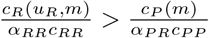, *then* 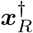 *is locally asymptotically stable*.

*Proof*. According to (25), if *c*_*R*_(*u*_*R*_, *m*) *>* 0, then *λ*_2_ *<* 0.

By dividing (24) and (26) by *α*_*SR*_*c*_*SS*_ and *α*_*PR*_*c*_*PP*_ respectively, we can see that *λ*_1_ and *λ*_3_ are negative when 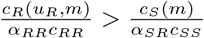, and 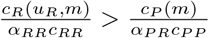. □

#### 3.3.1 Bifurcation analysis

All eigenvalues at the FRE are real, so a Hopf bifurcation is excluded. The FRE connects to the TE at *c*_*R*_(*u*_*R*_, *m*) = 0. A collision with CSR can occur at

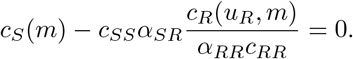

This boundary is independent of mutation. The persister invasion eigenvalue crosses zero with derivative

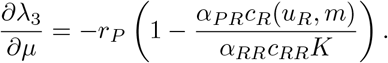

A nonzero value of (15) gives the generic transcritical collision with CRP.

### 3.4 Fully persister equilibrium

At this equilibrium, 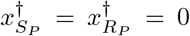, and we represent this equilibrium as 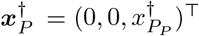. According to (2), from 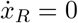 one can obtain:

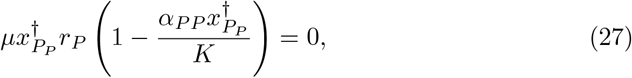

and from 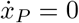 we have:

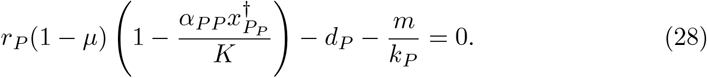

From (27) and (28), we can conclude the following:

1. 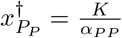 and dP = m = 0. These equations mean that for the existence of a fully persister equilibrium, there should be no killing agent, and persister cells should suppress apoptosis. With these conditions, the size of the tumor is close to the carrying capacity, which is not tolerable for the patient. Thus, a fully persister equilibrium does not exist or cannot be reached. As a result, stability analysis is not needed in this case.
2. 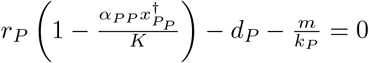
In this case, the mutation rate is zero. As we are interested in non-zero mutation rates, we do not consider this case here.

#### 3.4.1 Bifurcation analysis

For *d*_*P*_ + *m/k*_*P*_ *>* 0, there is no FPE branch and hence no FPE bifurcation in the assumed mutation interval. In the exceptional case *d*_*P*_ = *m* = 0, the population is independent of *µ*. If the resistant invasion expression above is negative, the quadratic has positive coefficients for every 0 *< µ <* 1; if it is positive, its constant coefficient is negative, and one eigenvalue is positive. Thus mutation alone produces no interior stability change or Hopf bifurcation at the FPE.

### 3.5 Coexistence of sensitive and resistant

We denote this equilibrium by 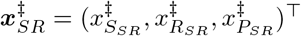, in which 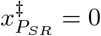. From 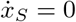 and 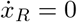, we can obtain

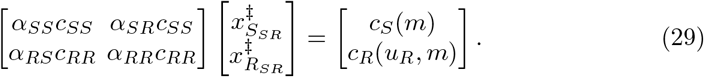

By defining

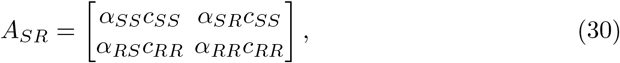

we can see that all elements of matrix (30) are positive. In addition, since 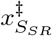 and 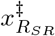 should be positive, we can conclude from (29) that *c*_*S*_(*m*) *>* 0 and *c*_*R*_(*u*_*R*_, *m*) *>* 0.

If det(*A*_*SR*_) *≠* 0, the equilibrium can be calculated as follows.

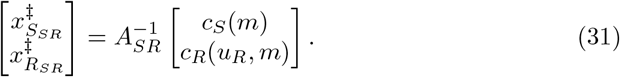

By using minors of the competition matrix, we can see that det 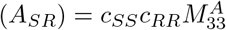.

For 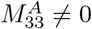, a CSR equilibrium exists if and only if both populations are positive:

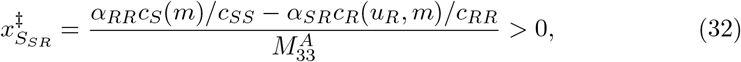

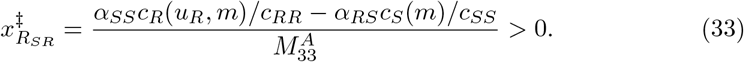

These inequalities imply *c*_*S*_(*m*) *>* 0 and *c*_*R*_(*u*_*R*_, *m*) *>* 0. Positivity of these two coef-ficients alone is not sufficient. If 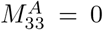, the two equations are compatible only when

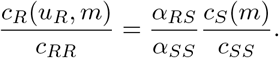

If compatibility holds and positive solutions exist, they form a line segment of equilibria. Otherwise there is no positive CSR equilibrium.

Substituting 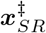 in the Jacobian (3), yields

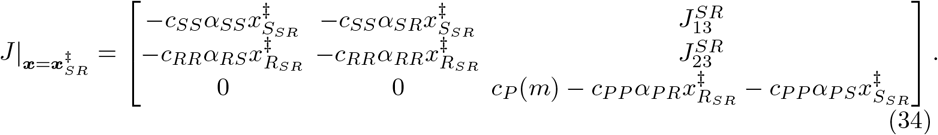

Thus, one of the eigenvalues is

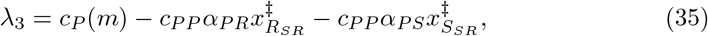

and the other two eigenvalues are the roots of the following quadratic equation. We can see that 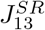 and 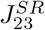 have no effect on the eigenvalues.

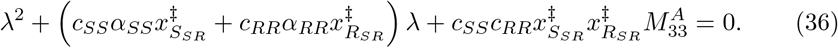

Consequently,

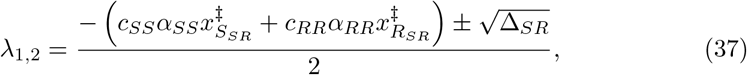

in which

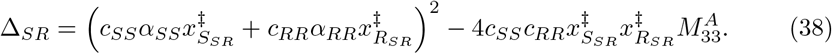

Since *c*_*SS*_ *>* 0 and *c*_*RR*_ *>* 0, having *λ*_1,2_ with negative real parts, requires 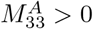.

#### Theorem 4.

*If* 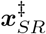 *exists, and additionally* 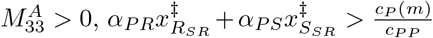*then* 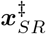 *is locally asymptotically stable*.

*Proof*. According to (37) and (38), if 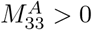, then *λ*_1,2_ will be negative or will have negative real parts.

By dividing (35) by *c*_*PP*_, we can see that if 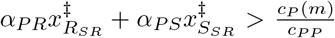, then *λ*_3_ *<* 0. □

#### Remark 2.

*According to Theorems 2, 3, for having a fully sensitive equilibrium and a fully resistant equilibrium, the following inequalities should hold:*

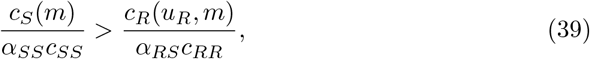

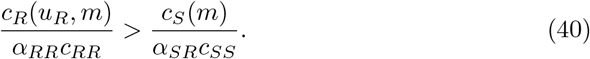

*These two conditions together result in*

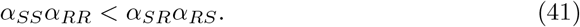

*Since* 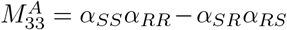 *follows directly from the definition of the* (3, 3) *minor of A, and consequently α*_*SS*_*α*_*RR*_ *< α*_*SR*_*α*_*RS*_ *is equivalent to* 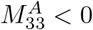 *definition. But for having a stable mixed equilibrium of resistant and sensitive cells* 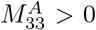. *Thus, if there exists a fully sensitive equilibrium and a fully resistant equilibrium, then it is not possible to have a stable mixed equilibrium of resistant and sensitive cells*.

#### 3.5.1 Bifurcation analysis

At a positive CSR equilibrium, the sensitive and resistant eigenvalues, *λ*_1,2_ are real. The discriminant in (38) can be rewritten as

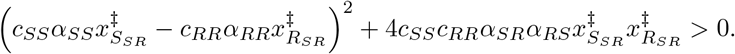

The trace of the 2 ×2 Jacobian of the sensitive-resistant matrix is negative. For 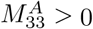, both are negative; for 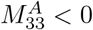, one is positive. Consequently, CSR cannot undergo a Hopf bifurcation. If 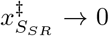 while the resistant population remains positive, the CSR branch reaches the fully resistant equilibrium. 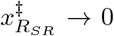 while the sensitive population remains positive, it reaches the fully sensitive equilibrium. Moreover, the compatible case 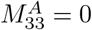 gives a continuum rather than an isolated saddle-node.

The persister invasion eigenvalue is

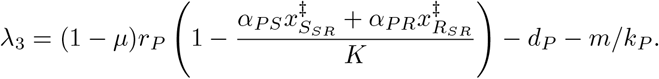

The sensitive and resistant populations do not depend on mutation. At a mutation value strictly between zero and one where the persister-invasion eigenvalue vanishes, the derivative with respect to *µ* is negative. With a nonzero coefficient in (15), this gives a transcritical collision with CAT.

### 3.6 Coexistence of sensitive and persister cells

At this equilibrium which we represent by 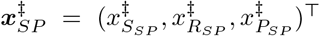, we have 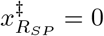. From 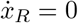 in (2), we can obtain:

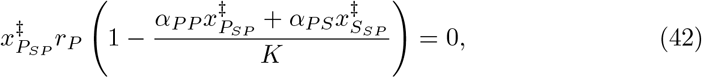

which yields

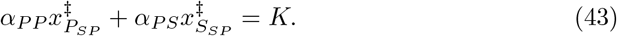

In addition, from 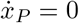, we have

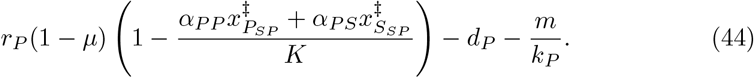

From (43) and (44), one can obtain

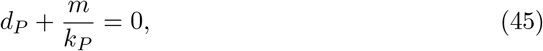

which means *d*_*P*_ = *m* = 0. We can see that in a special case in which *d*_*P*_ = 0 and there exists no treatment, we can have a coexistence of sensitive and persister cells.

But, according to (43), if this equilibrium exists and is stable, tumor will be stabilized on a volume which is not tolerable for the patient. Thus, tumor burden is very high.

We do not add stability analysis for this case because this equilibrium is not reachable.

### 3.7 Coexistence of resistant and persister cells

At this equilibrium, 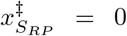, and we represent this equilibrium as 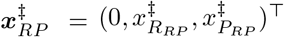. According to (2), from 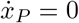 one can obtain:

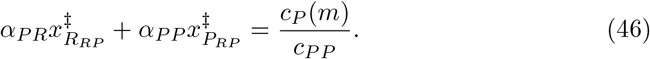

and from 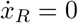 in (2), we have

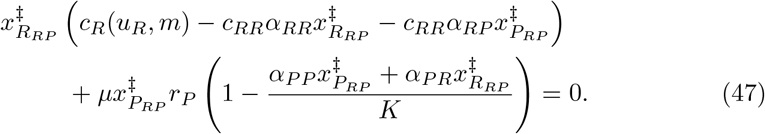

By substituting (46) in (47),

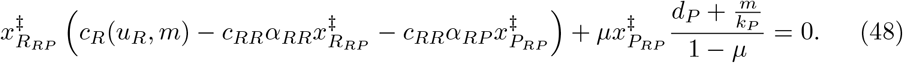

After some algebraic manipulations, using minors of the competition matrix, and by substituting 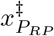 as a function of 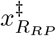 from (46) in (48), we have the following quadratic equation with respect to 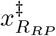.

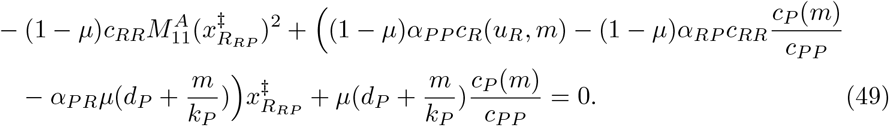

By defining the following matrix

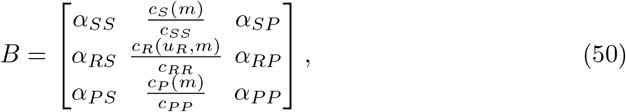

the coefficient of 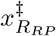 in (49) can be reduced by using minors of the matrix (50):

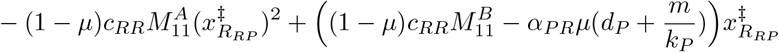

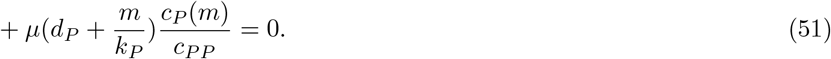

The discriminant Δ_*RP*_ of (51) is defined as follows.

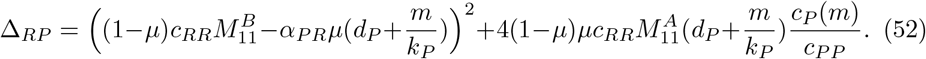

If 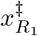 and 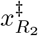 are the roots of (51), we have the following relations between the roots.

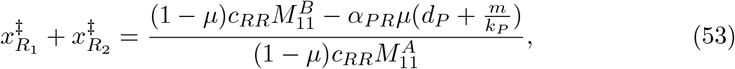

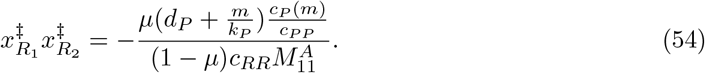

The root relations above require 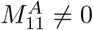. Every CRP equilibrium must satisfy

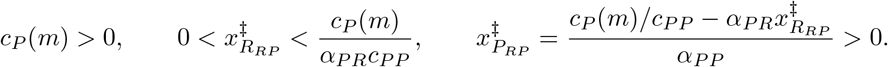

For 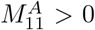, the quadratic has one positive and one negative root. Its positive root is biologically acceptable only if it satisfies the upper bound that is tolerable for the patient. This is a sufficient condition for one positive root, and is not a necessary condition for a unique biologically acceptable equilibrium. For 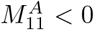, two positive roots require

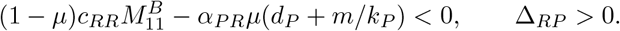

Both roots must still satisfy the population bounds. If 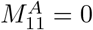, equation (51) is linear; its root is obtained by dividing minus the constant term by the linear coefficient when that coefficient is nonzero. For *c*_*P*_ (*m*) *>* 0 and *d*_*P*_ + *m/k*_*P*_ *>* 0, a zero linear coefficient gives no root. A zero discriminant yields a repeated root, whose biological relevance must be checked.

Substituting 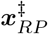 in the Jacobian (3), yields

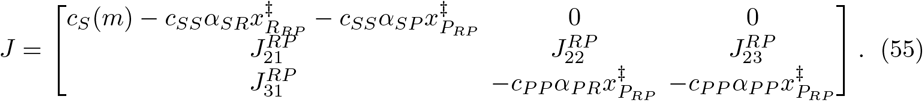

in which

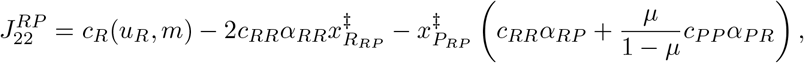

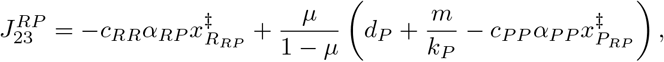

and we should mention that 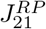, and 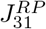 have no effects on the eigenvalues. The first eigenvalue of (55) is

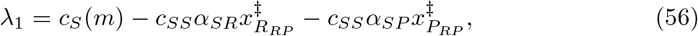

and the other two eigenvalues are the roots of the following equation.

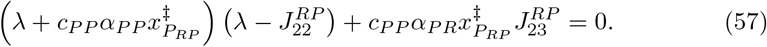

By algebraic manipulations, (57) can be rewritten as follows.

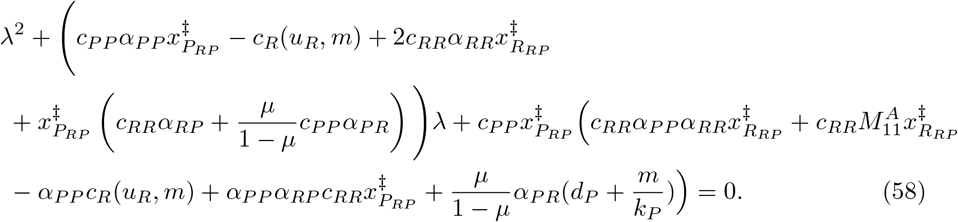

#### Theorem 5.

*If* 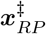 *exists, and additionally*

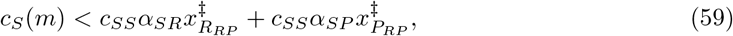

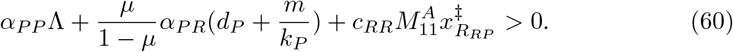

*in which*

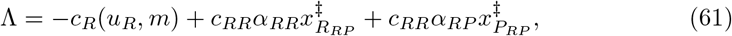

*then* 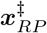 *is locally asymptotically stable*.

*Proof*. According to (56), if 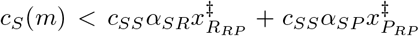, then *λ*_1_ will be negative.

Assume that *λ*_2_ and *λ*_3_ are the roots of (57). Then, we have the following relations between the roots.

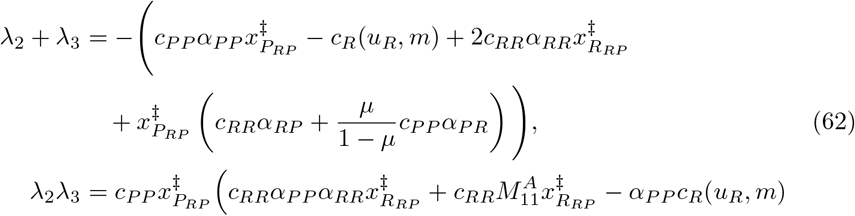

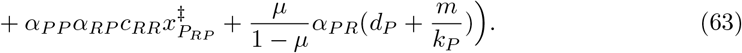

According to (48), we have

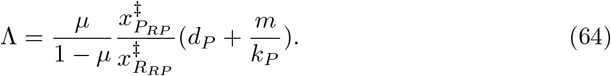

Since the right-hand side of (64) is positive, we can conclude that Λ*>* 0. Moreover, (62) and (63) can be rewritten as follows:

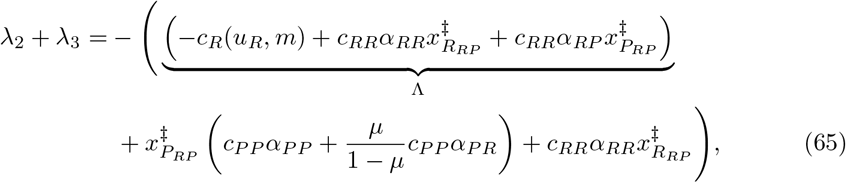

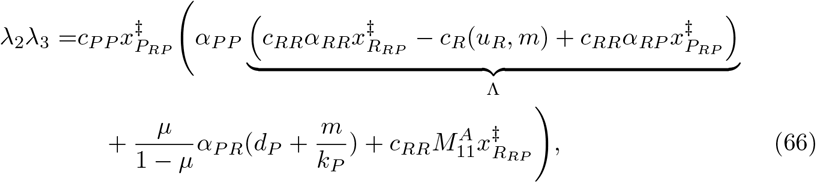

since Λ*>* 0, we can conclude that *λ*_2_ + *λ*_3_ *<* 0. In this situation, three scenarios are possible: 1) both *λ*_2_ and *λ*_3_ are real negatives, 2) *λ*_2_ and *λ*_3_ are complex conjugates with negative real parts, 3) one of the *λ*_2_ and *λ*_3_ is negative and the other is positive. The only scenario that will result in an unstable eigenvalue is the third scenario. In the third scenario, *λ*_2_*λ*_3_ *<* 0, while in the first two scenarios *λ*_2_*λ*_3_ *>* 0. Thus, if *λ*_2_*λ*_3_ *>* 0, we will have stable eigenvalues. For *λ*_2_*λ*_3_ *>* 0,, we should have

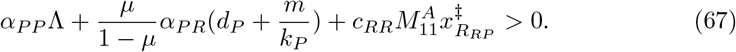

Since Λ*>* 0, if 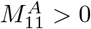, then *λ*_2_*λ*_3_ *>* 0, and *λ*_2_ and *λ*_3_ are stable eigenvalues. Thus, if among the resistant and persister cells, competition between the same types is stronger than the competition between different types, then *λ*_2_ and *λ*_3_ have negative real parts. Thus, *α*_*RR*_*α*_*PP*_ *> α*_*RP*_ *α*_*PR*_ is sufficient for these two eigenvalues to have negative real parts. This condition compares products of competition coefficients. □

#### 3.7.1 Bifurcation analysis

For a positive CRP equilibrium and *d*_*P*_ + *m/k*_*P*_ *>* 0, equation (64) gives Λ*>* 0. Hence

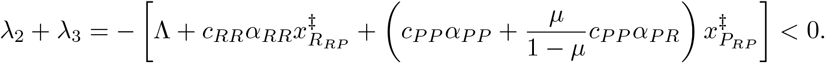

The resistant–persister pair therefore cannot produce a Hopf bifurcation. The remaining eigenvalue is real, so a Hopf bifurcation is excluded for CRP.

A saddle-node candidate occurs when Δ_*RP*_ = 0 and 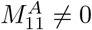. The repeated root is

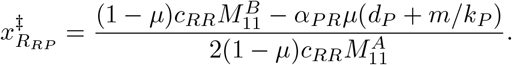

The populations must satisfy the strict bounds above. The quadratic coefficient is nonzero under these assumptions. A nonzero derivative of the discriminant with respect to the varied parameter gives a simple fold of the equilibrium equation. The sensitive invasion eigenvalue must also be nonzero for a simple saddle-node in the full system.

The boundaries 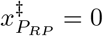 and *λ*_1_ = 0 connect CRP to FRE and CAT, respectively. At the latter boundary, the center direction is the sensitive population. The linear responses on the center manifold satisfy

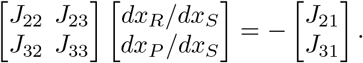

The quadratic coefficient is

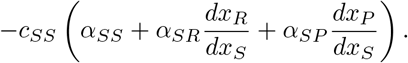

A nonzero coefficient and a nonzero derivative of the sensitive eigenvalue along the CRP branch establish the generic transcritical case.

### 3.8 Coexistence of all three types of cancer cells

From 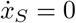, and 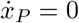, we can obtain the following.

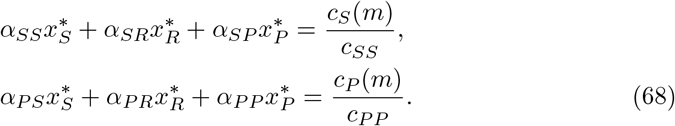

From 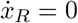, we have

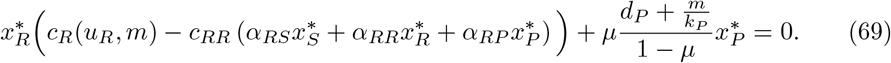

For 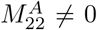, using equations (68), we can rewrite 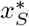 and 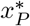 as functions of 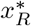 as follows.

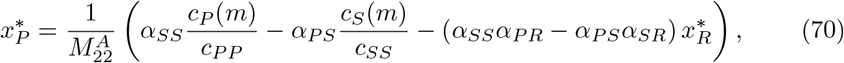

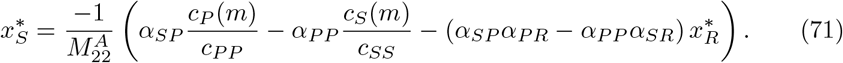

By substituting 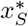 and 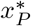 from (71) and (70) into (69), and after some manipulations, we obtain a quadratic equation with respect to 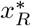:

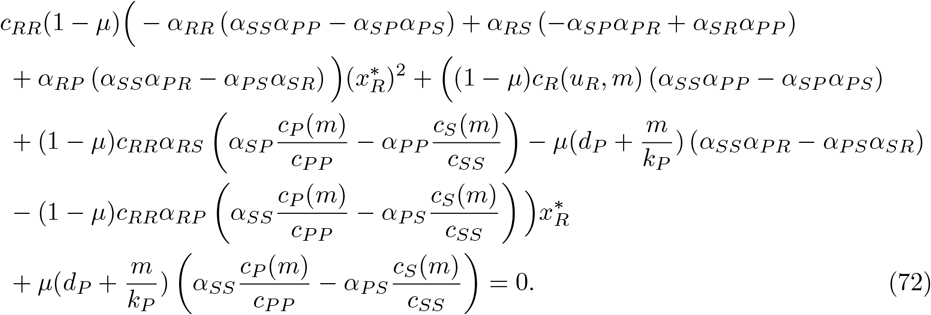

Using minors of the competition matrix, the coefficient of 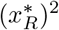 in (72) can be rewritten as follows.

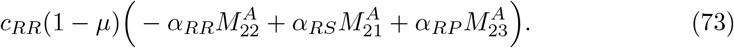

But according to the Laplace expansion of the determinant of the competition matrix on its second row, the coefficient of 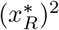 in (72) can be reduced to *−c*_*RR*_(1*−µ*)det(*A*).

Moreover, we can simplify the coefficient of 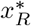 in (72), using minors of the competition matrix as follows.

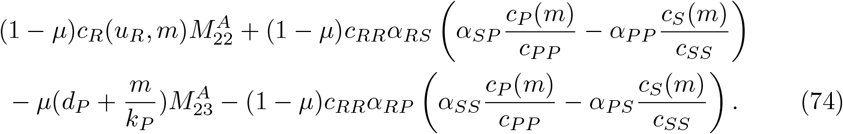

According to the matrix B (50), the coefficient of 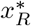 in (72) becomes

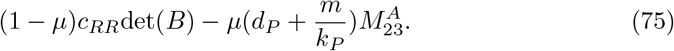

Thus, the quadratic equation (72) reduces to

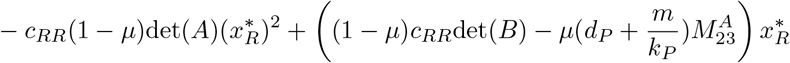

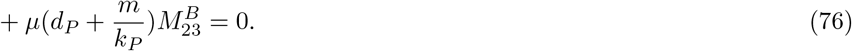

For det(*A*) ≠ 0, 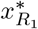 and 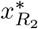 are the two solutions for the quadratic equation (76), we know

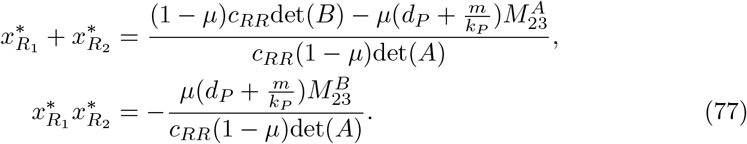

The signs of the root sum and product characterize algebraic roots, but are not sufficient for biological coexistence. Every accepted root must be real and satisfy 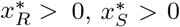, and 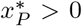 after substitution in (70)-(71). In particular, *c*_*S*_(*m*) *>* 0 and *c*_*P*_ (*m*) *>* 0 are necessary. Define the equilibrium discriminant by

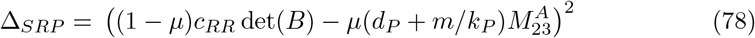

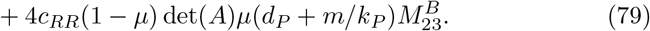

One positive and one negative algebraic root occur when det 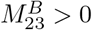. Two distinct positive algebraic roots require 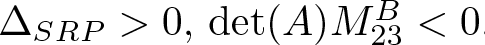, and

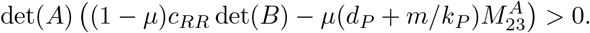

In either case, all population inequalities must also hold. If det(*A*) = 0, equation (76) is linear or degenerate. If 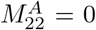, equations (68) and (69) must be solved directly, using another nonzero minor when possible.

#### Theorem 6.

*If* 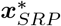 *exists, and additionally*

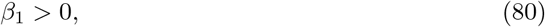

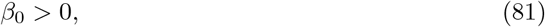

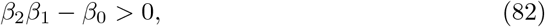

*in which β*_*i*_ *is the coefficient of λ*^*i*^ *in the following cubic equation for i ∈ {*0, 1, 2*}*,

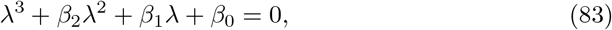

*where*

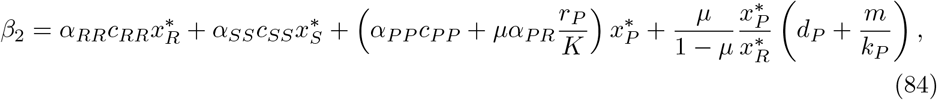

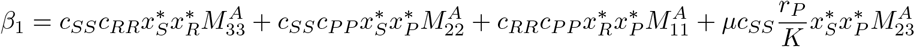

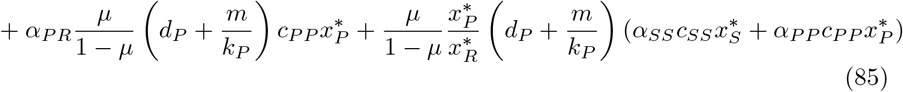

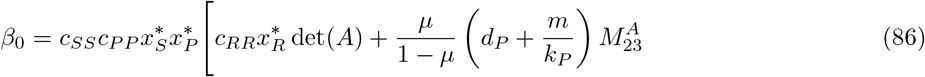

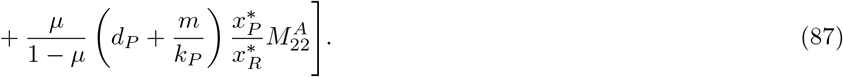

*then* 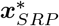 *is locally asymptotically stable*.

*Proof*. For checking the stability, we should substitute the 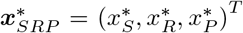 in the Jacobian matrix (3).

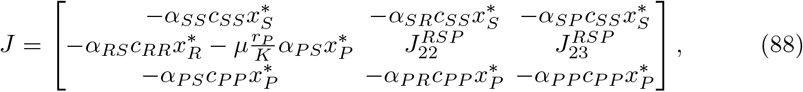

in which

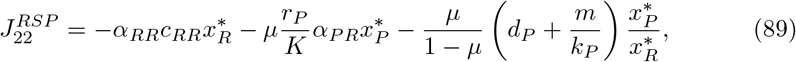

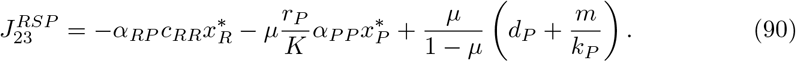

The eigenvalues of this Jacobian matrix are the roots of the cubic equation (83). The constant terms in (83) can be simplified by using minors of the competition matrix.

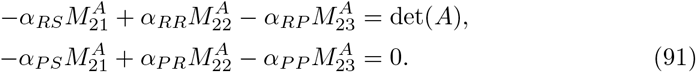

At locally stable mixed equilibria, all roots of (83) have negative real parts. If *λ*_1_, *λ*_2_, and *λ*_3_ are the roots of (83), they satisfy:

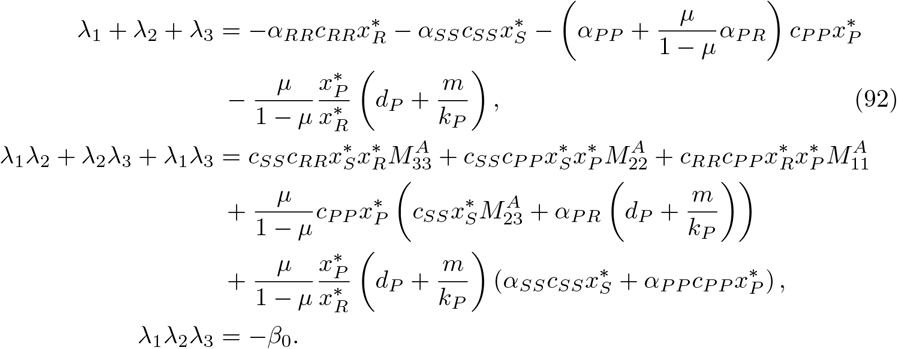

By defining *β*_*i*_ as the coefficient of *λ*^*i*^ in (83) for *i ∈ {*0, 1, 2}, and applying the Routh–Hurwitz criterion [26], we have the following conditions for having a locally stable equilibrium:

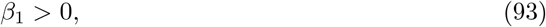

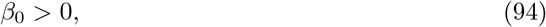

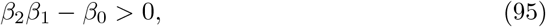

Since *µ ∈* (0, 1), all the parameters on the right-hand side of (92) are negative and we can conclude that *λ*_1_ + *λ*_2_ + *λ*_3_ *<* 0, and as a result, *β*_2_ *>* 0. □

It should be mentioned that in a special case, the condition *β*_2_*β*_1_ *− β*_0_ *>* 0 for having a locally stable equilibrium can be simplified. According to the fundamental theorem of algebra, (83) can have three real roots or one real root and two complex conjugate roots. By introducing the following variables

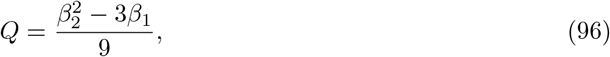

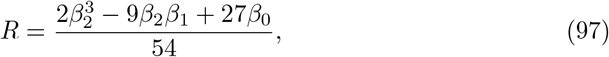

we use *R*^2^ *− Q*^3^ to classify the eigenvalues. This expression is different from the equilibrium discriminant Δ_*SRP*_. If *R*^2^ *−Q*^3^ *<* 0, all three eigenvalues are real; if *R*^2^ *−Q*^3^ *>* 0, one is real and two are complex conjugates. Equality gives repeated real roots.

When all eigenvalues are real, *β*_2_ *>* 0, *β*_1_ *>* 0, and *β*_0_ *>* 0 imply that they are negative. Indeed, suppose that *λ*_1_ *<* 0 and *λ*_2_, *λ*_3_ *>* 0. The negative sum and positive pairwise sum imply

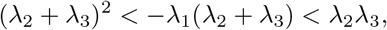

and hence

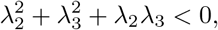

which is impossible. Thus, under the additional condition *R*^2^ *− Q*^3^ *≤* 0, *β*_1_ *>* 0 and *β*_0_ *>* 0 suffice. In the general case, all Routh–Hurwitz inequalities must be checked.

#### 3.8.1 Bifurcation analysis

Direct calculation gives *β*_0_ = *−* det(*J*) and *β*_2_ = *−* tr(*J*) *>* 0. A saddle-node candidate of two CAT branches requires Δ_*SRP*_ = 0, det(*A*) ≠ 0, and positive populations. For a simple fold, the varied parameter must cross the discriminant transversely, and the full Jacobian must have only one zero eigenvalue. At *β*_0_ = 0, the latter requires *β*_1_≠0. A fold does not follow from the eigenvalue discriminant *R*^2^*− Q*^3^ = 0, which only indicates repeated eigenvalues.

Transcritical collisions occur when a CAT population reaches zero, and the neighboring equilibrium exists. Under positive persister mortality, 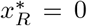 with 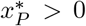 is impossible. Hence the relevant boundaries connect CAT to CRP at 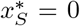, or to a persister-free equilibrium at 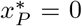. The center-manifold calculations given above distinguish a nondegenerate crossing from a higher-order case.

An interior Hopf candidate satisfies

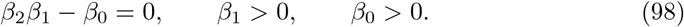

At this point the characteristic polynomial factors as

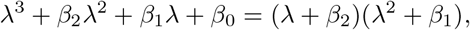

so the eigenvalues are *−β*_2_ and 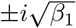. Differentiation of the characteristic equation along the equilibrium branch gives

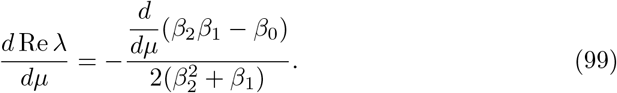

The derivative in (99) includes the changes in the equilibrium populations as mutation varies. We keep *m* and all other parameters fixed and differentiate the three equilibrium equations 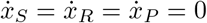 from (2) with respect to *µ*. The population derivatives are then obtained by solving the linear system

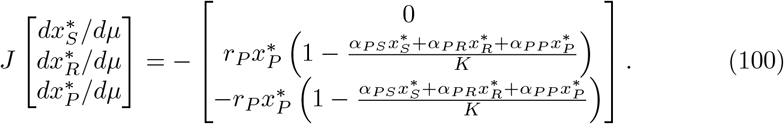

All quantities are evaluated at the equilibrium and mutation value under consideration. At a Hopf point, *J* has no zero eigenvalue, so this system has a unique solution. We use these population derivatives when differentiating the characteristic coefficients:

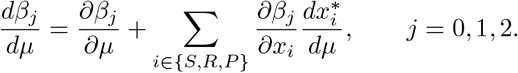

The partial derivatives are taken before substituting the equilibrium populations. Substituting the resulting total derivatives into (99) gives the rate at which the real part of the critical eigenvalue changes with mutation. A nonzero rate verifies transversality.

## 4 Illustration of cancer trajectories, equilibrium distributions, and bifurcations

In this section, we show the impact of different treatment doses and mutation rates on the stability of different equilibria. We illustrate under which conditions we have the coexistence of different equilibria of the system.

Parameter values used for the simulations are listed in Table 2. Moreover, unless specified otherwise, we use *u*_*R*_ = 3.3 and

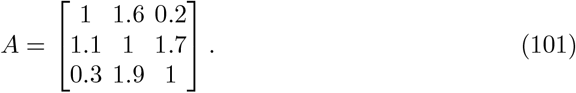

### 4.1 Distribution of different equilibria

Here, we illustrate how different equilibria are distributed in the (*µ, m*)- and (*u*_*R*_, *m*)-planes.

Distribution of equilibria in the (*µ, m*)-plane for a given set of parameters is demonstrated in Fig. 1. We can see that, for high mutation rates, most stable equilibria have one or zero cell types, such as FRE, FSE, and TE. Moreover, from Fig. 1, it is demonstrated that for having more than one stable equilibrium, the treatment dose should be low enough. For instance, we can observe that for *m <* 0.4, we can have two stable FSE and FRE, or FRE and CAT. Thus, for *m <* 0.4,, depending on the initial tumor composition and basins of attraction of the stable equilibria, we can stabilize tumor burden on one of the equilibria.

**Fig. 1:**
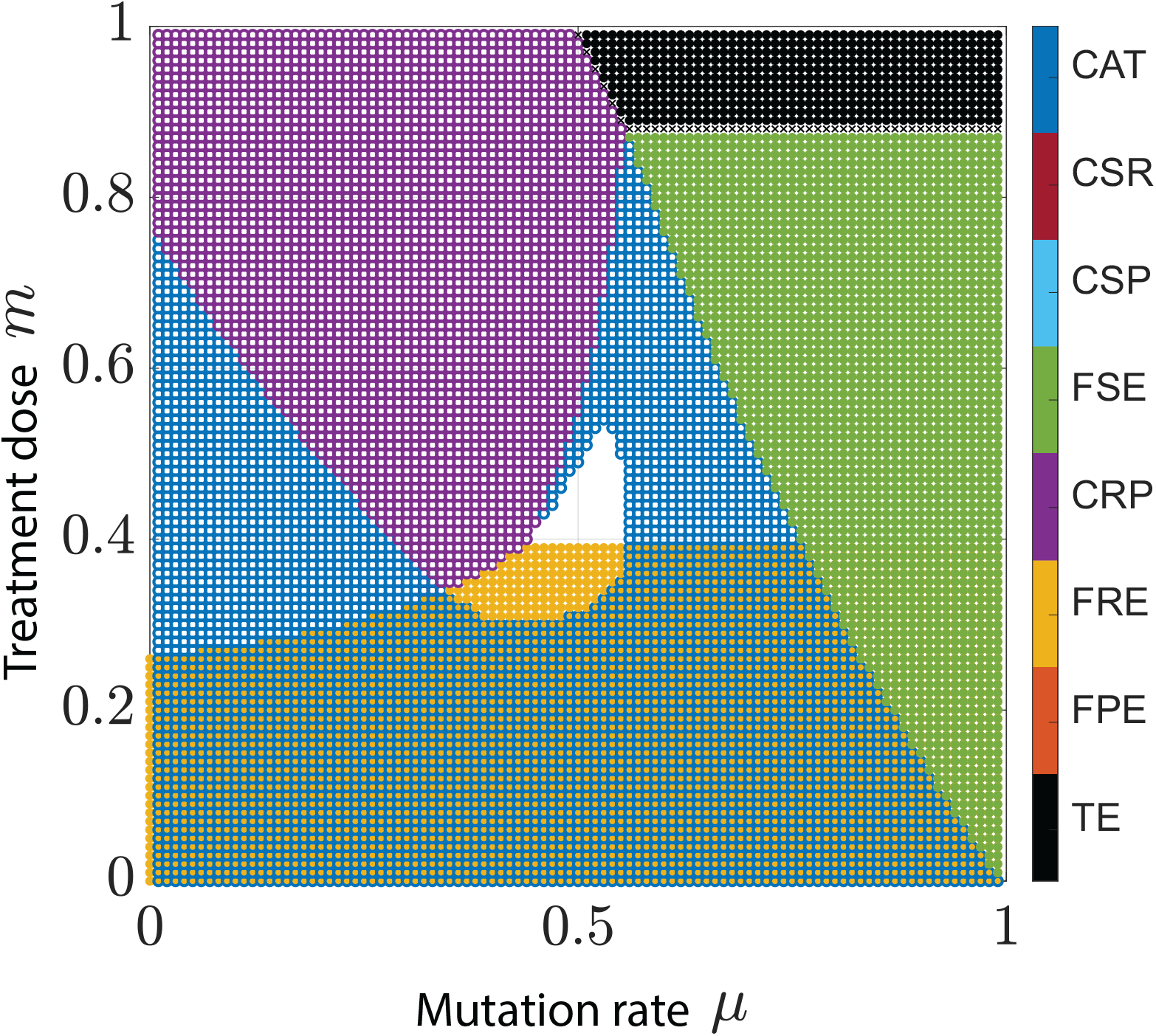
Distribution of equilibria in the (*µ, m*)-plane. For the parameters in the white area, there exists no stable equilibria. Parameters: *u*_*R*_ = 3.3, *α*_*SR*_ = 1.6, *α*_*SP*_ = 0.2, *α*_*RS*_ = 1.1, *α*_*RP*_ = 1.7, *α*_*PR*_ = 1.9, *α*_*PS*_ = 0.3. TE: Trivial Equilibrium, FPE: Fully Persister Equilibrium, FRE: Fully Resistant Equilibrium, FSE: Fully Sensitive Equilibrium, CRP: Coexistence of Resistant and Persister, CSP: Coexistence of Sensitive and Persister, CSR: Coexistence of Sensitive and Resistant, and CAT: Coexistence of All Three types of cells. There will be no stable equilibrium for the (*µ, m*) in the white region.

Moreover, in Fig. 1, the equilibria with the zero eigenvalues are demonstrated by crosses “x”. For instance, we can see these equilibria in Fig. 1, demonstrating the transition from the green (FSE) to the black (TE) region.

We can see from Fig. 1, that for a fixed treatment dose, with higher mutation rates the parameter region in which the FSE is locally stable is increased. Moreover, for the existence of a locally stable TE, both the treatment dose and mutation rate should be high enough.

In the sequel, we consider the case in which *µ* is a function of treatment dose *m*. In the real world, when there exists no treatment, the mutation rate is high. While in the presence of high treatment doses, the mutation rate is low and negligible. Consequently, we assume that the relation between the mutation rate and treatment dose is as follows. By assuming that *u*_*R*_ *∈* [0, 10], the distribution of equilibria in the (*u*_*R*_, *m*)-plane is demonstrated in Fig. 2.

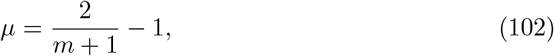

**Fig. 2:**
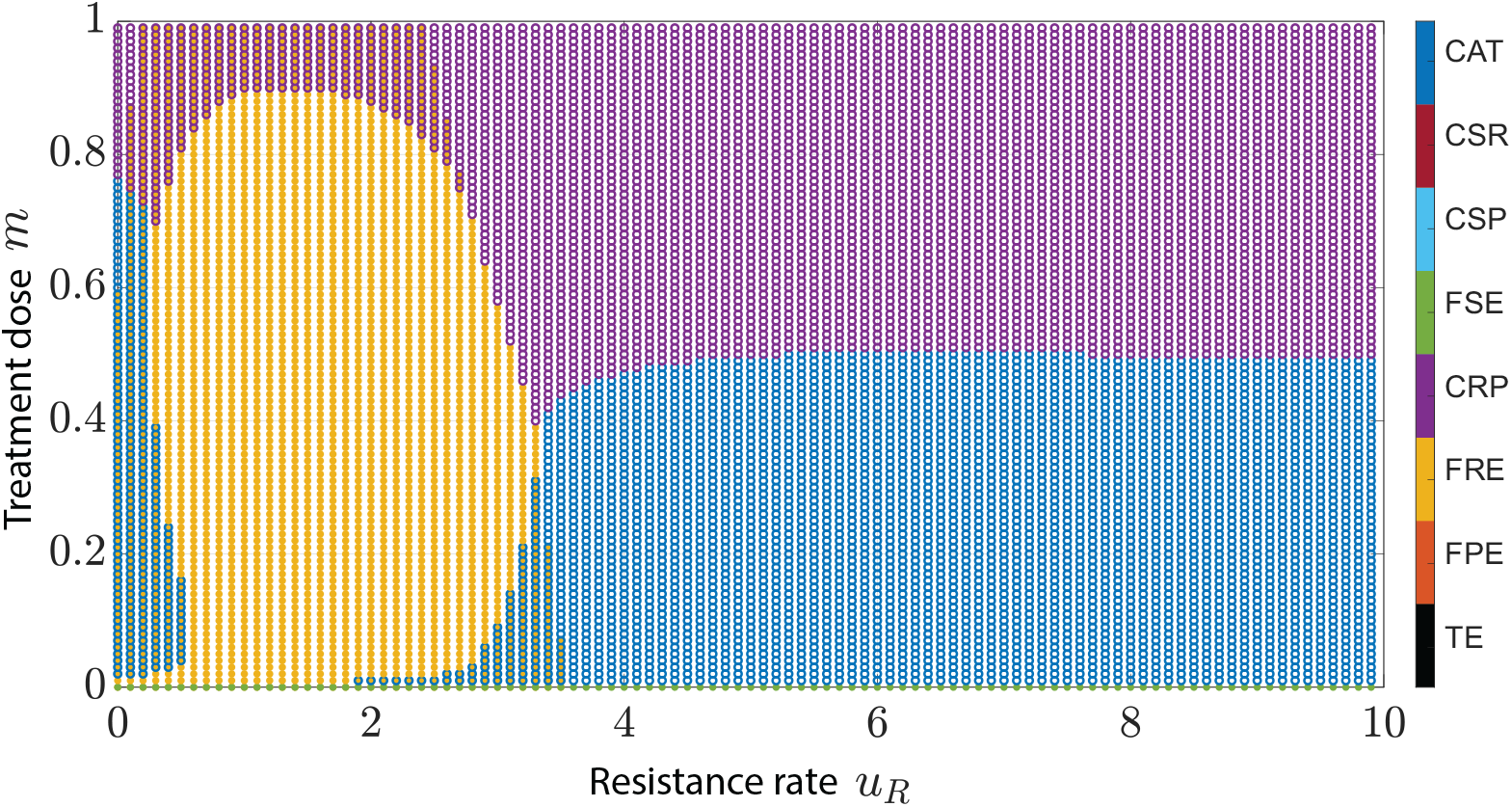
Distribution of different equilibria in the (*u*_*R*_, *m*)-plane when the mutation rate is a decreasing function of the treatment dose. Parameters: *α*_*SR*_ = 1.6, *α*_*SP*_ = 0.2, *α*_*RS*_ = 1.1, *α*_*RP*_ = 1.7, *α*_*PR*_ = 1.9, *α*_*PS*_ = 0.3. TE: Trivial Equilibrium, FPE: Fully Persister Equilibrium, FRE: Fully Resistant Equilibrium, FSE: Fully Sensitive Equilibrium, CRP: Coexistence of Resistant and Persister, CSP: Coexistence of Sensitive and Persister, CSR: Coexistence of Sensitive and Resistant, and CAT: Coexistence of All Three types of cells.

We can see from Fig. 2, that for *u*_*R*_ *>* 4, only two stable equilibria can be expected. If *m* is high enough, there exists one stable CSR, while for low treatment doses, CAT is the only stable equilibrium. Moreover, it can be observed from Fig. 2, that a stable fully resistant equilibrium (FRE) can be expected only for *u*_*R*_ *<* 4.

### 4.2 Numerical bifurcation analysis

For *m* = 0.45, Table 3 reports every biologically feasible equilibrium collision and Hopf bifurcation in 0 *< µ <* 1 for the competition matrix (101). The values are calculated from invasion eigenvalues, the equilibrium discriminants, and the Hopf condition (98). Substitution into the original equations verifies the candidate populations. Figure 3 shows both stable and unstable feasible branches.

**Table 3:** Bifurcation values for the given parameter set at *m* = 0.45.

| $\mu$ | Equilibria and type | Result as mutation increases |
| --- | --- | --- |
| 0.241 | CAT–CRP, transcritical | Stable CAT is replaced by stable CRP. |
| 0.465 | CRP–CAT, transcritical | Stable CRP is replaced by stable CAT. |
| 0.477 | CAT, Hopf | CAT loses stability. |
| 0.492 | CRP–FRE, transcritical | Both branches are unstable to sensitive invasion. |
| 0.557 | CAT, Hopf | CAT becomes stable. |
| 0.732 | CAT–FSE, transcritical | Stable CAT is replaced by stable FSE. |

**Fig. 3:**
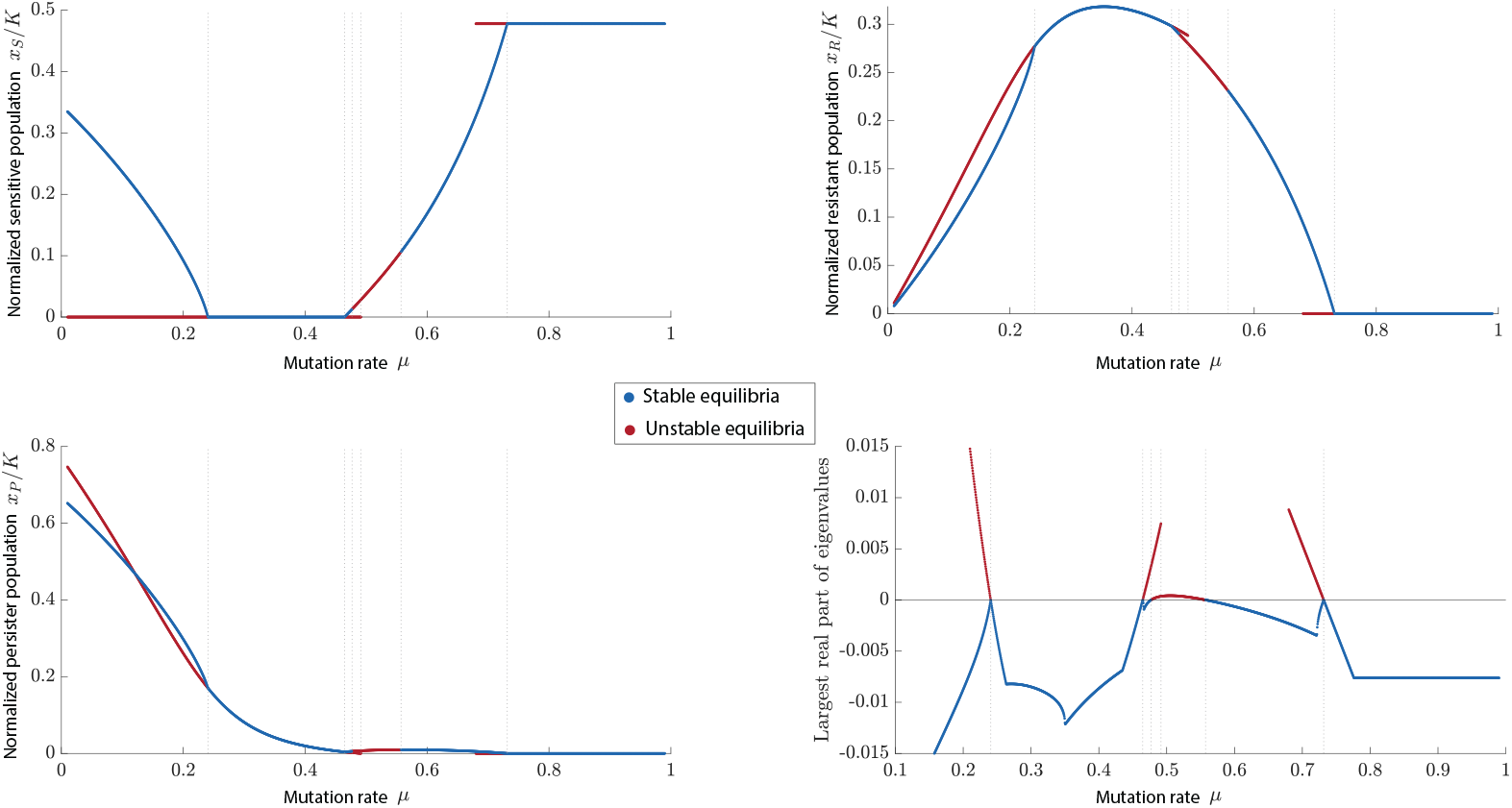
Feasible equilibrium branches at *m* = 0.450. Blue points indicate strictly stable equilibria and red points indicate unstable equilibria. Dotted vertical lines mark the six values in Table 3. The TE and FRE, and the unstable FSE below *µ* = 0.680, are omitted to keep the coexistence branches visible. The bottom right figure displays the largest real part of the eigenvalues, with the vertical scale restricted to the neighborhood of zero.

The stable intervals are CAT on (0, 0.241), CRP on (0.241, 0.465), CAT on (0.465, 0.477), no stable equilibrium on (0.477, 0.557), CAT on (0.557, 0.732), and FSE on (0.732, 1). The FRE–CRP collision inside the interval without a stable equilibrium does not change this list. The TE persister eigenvalue is zero at *µ* = 0.77, but the adjacent CRP branch is infeasible, and the TE remains unstable.

At the first Hopf point, 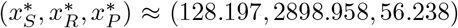 and the eigen-values are *−* 0.017 and *±*0.003*i*. At the second, the populations are approximately (1063.574, 2309.166, 97.689) and the *ω* is 0.007. The rates of changes in the real part of the critical eigenvalue with respect to *µ* in (99) are 0.0449 and *−* 0.013, respectively, as shown in Fig. 4.

**Fig. 4:**
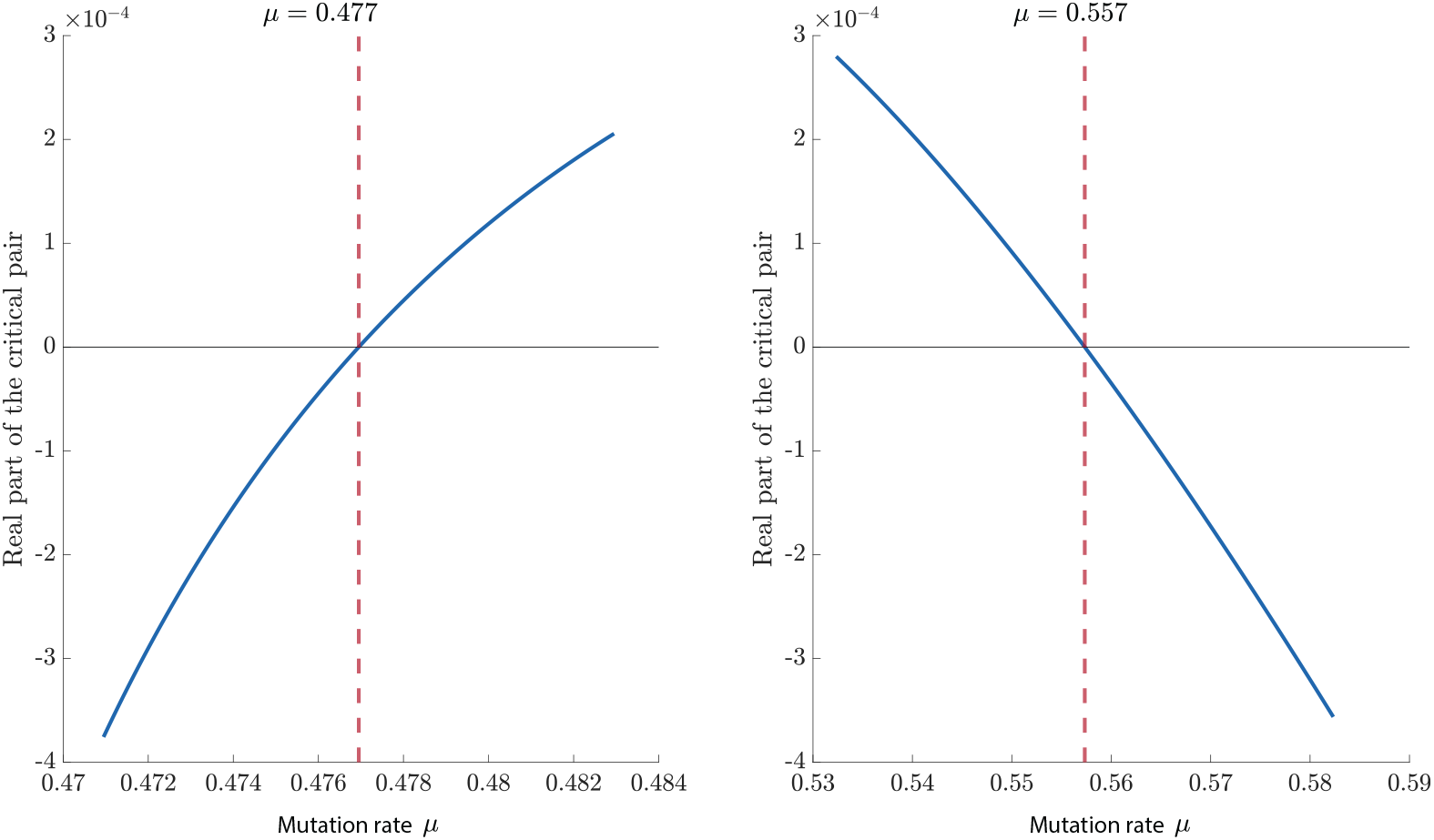
The critical complex eigenvalue pair crosses the imaginary axis at *µ* = 0.477 and *µ* = 0.557, with *m* = 0.450. The dashed lines give the calculated Hopf locations. The other eigenvalue is negative at both points.

**Fig. 5:**
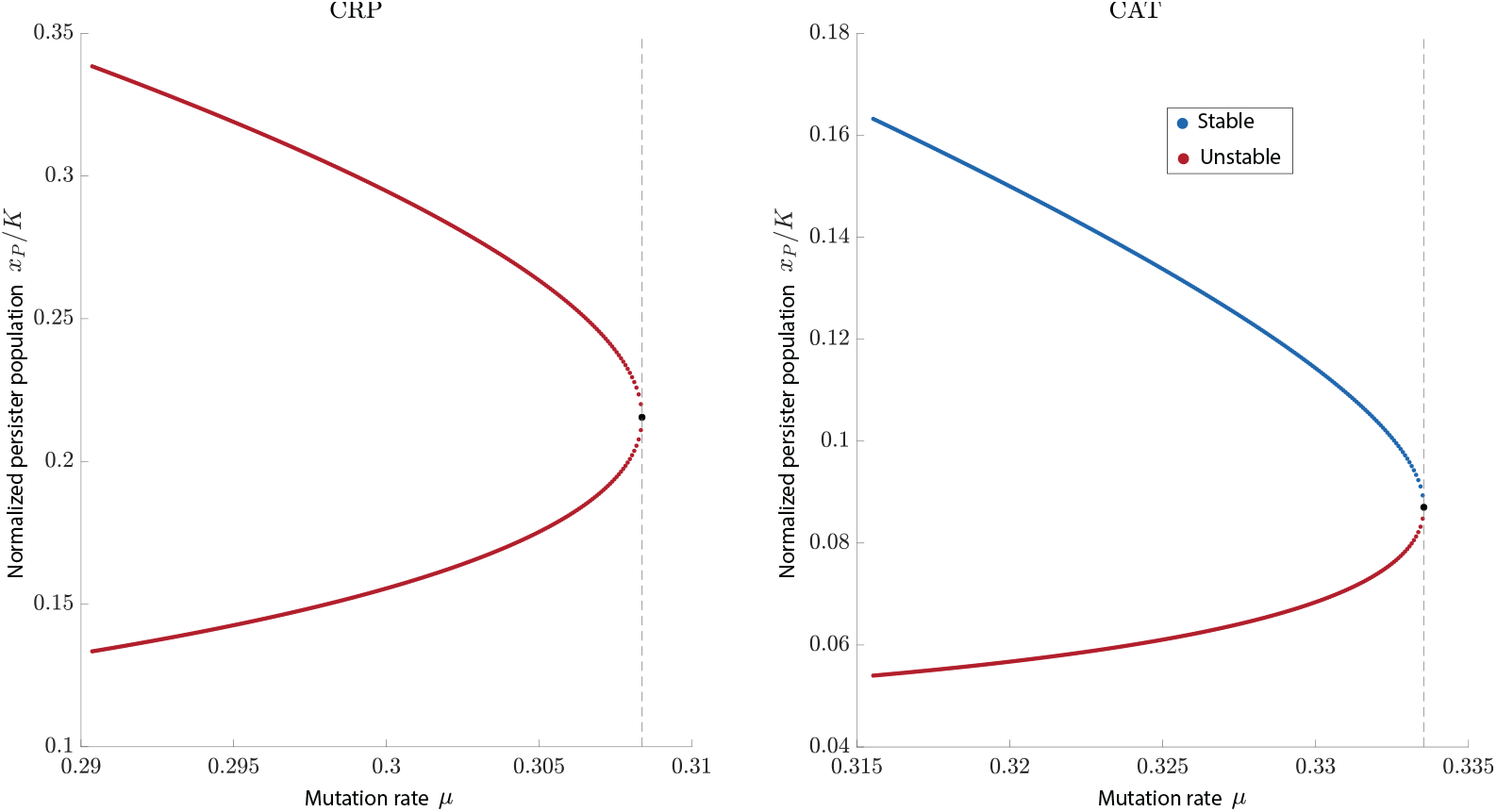
Positive saddle-node bifurcations. Left: CRP at (*m, µ*) = (0.200, 0.308). Right: CAT at (*m, µ*) = (0.350, 0.334). Blue denotes stable branches, red denotes unstable branches, and black circles mark the folds. The left pair is unstable to sensitive invasion on both branches.

The reported eigenvalue crossing rates are obtained by solving (100) at each Hopf equilibrium and substituting the resulting population derivatives into (99). The first Lyapunov coefficients are calculated separately from the Jacobian and the second derivatives of the right-hand sides of (2), evaluated at the same equilibria. The original equations are quadratic in the populations, so their third derivatives are zero. This calculation uses the original population variables and does not require their derivatives with respect to *µ*.

At both Hopf points, the nonlinear analysis predicts small stable oscillations of the sensitive, resistant, and persister populations. The calculated first Lyapunov coefficients are negative, approximately *−*5.134 *×* 10^*−*6^ and *−*2.120 *×* 10^*−*8^, when the coefficients are calculated using the original population variables *x*_*S*_, *x*_*R*_, and *x*_*P*_, without dividing them by *K*. Therefore, both Hopf bifurcations are supercritical. Small stable periodic solutions exist for mutation rates slightly above 0.477 and slightly below 0.557, where the CAT equilibrium is unstable. These conclusions apply near the two bifurcation points. These calculations do not establish whether the periodic solutions near the two Hopf points belong to the same continuous branch for mutation rates between 0.477 and 0.557.

The calculations also confirm transcritical bifurcations where two equilibrium branches meet. At mutation rates *µ ≈*0.241 and *µ ≈*0.465, the CRP and CAT branches meet. The eigenvalue that determines whether a small sensitive population grows or declines near CRP changes sign at these points. Its rates of change with respect to mutation are approximately *−* 0.416 and 0.255, respectively. The corresponding coefficients of the quadratic terms, which describe the leading nonlinear effects near these intersections, are *−*1.07818 *×*10^*−*5^ and*−* 2.47186 *×*10^*−*5^.

At the FRE–CRP and FSE–CAT intersections, the relevant eigenvalue determines whether a small persister population grows or declines. Its rates of change with respect to mutation are approximately *−*0.090 and *−*0.171, respectively. The corresponding quadratic coefficients are *−*7.017 *×*10^*−*5^ and *−*1.304 *×*10^*−*4^. These coefficients are calculated using the original population variables, without dividing them by *K*. At each of the four intersections, exactly one eigenvalue is zero, its rate of change with mutation is nonzero, and the corresponding quadratic coefficient is nonzero. These results verify the conditions for the transcritical bifurcations.

To illustrate saddle-node bifurcations, we retain the same competition matrix and consider two treatment doses. For each dose, we vary the mutation rate. At *m* = 0.20, two CRP equilibria meet at *µ ≈* 0.308, where the populations are approximately 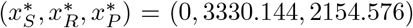. The eigenvalues are approximately 0, *−*0.072,and 0.081. The positive eigenvalue means that a small sensitive population introduced near this equilibrium initially grows. Consequently, both nearby CRP branches are unstable when changes in all three populations are considered. At *m* = 0.35, two CAT equilibria meet at *µ≈* 0.333537, where the populations are approximately (402.782, 3320.032, 870.280). One eigenvalue is zero, and the other two are approximately *−*0.046 and *−*0.010. In this case, one of the two nearby CAT branches is stable, whereas the other is unstable.

We also checked the mathematical conditions required for these saddle-node bifurcations. Near each meeting point, the behavior associated with the zero eigenvalue can be described by a simplified equation. This equation includes a term describing the effect of changing the mutation rate and a quadratic term describing the leading nonlinear effect of deviations from the equilibrium. For the CRP and CAT cases, respectively, the coefficients of the mutation-rate terms are approximately *−*101.544 and *−*58.066, and the quadratic coefficients are *−*1.384 *×*10^*−*6^ and 1*−*.847 *×*10^*−*6^. These values are calculated using the original population variables, without dividing them by *K*. At each meeting point, exactly one eigenvalue is zero and both coefficients are nonzero, confirming the conditions for a saddle-node bifurcation. Reversing the direction used to represent deviations in the simplified equation changes the signs of both coefficients but does not change this conclusion.

### 4.3 Illustration of typical cancer trajectories

Typical cancer trajectories that are attracted to some stable equilibria are demonstrated in this subsection.

Figure 6 compares three mutation values at *m* = 0.45, starting from (*x*_*S*_(0), *x*_*R*_(0), *x*_*P*_ (0)) = (1000, 1000, 1000). At *µ* = 0.45, the trajectory approaches CRP. At *µ* = 0.65, it approaches CAT. At *µ* = 0.50, the trajectory displays sustained oscillations over the simulated interval 0 *≤ t≤* 50000. The corresponding CAT has a complex eigenvalue pair with positive real part. The absence of a stable equilibrium alone does not establish oscillatory behavior. Here the direct simulations and the two local Hopf calculations provide separate evidence.

**Fig. 6:**
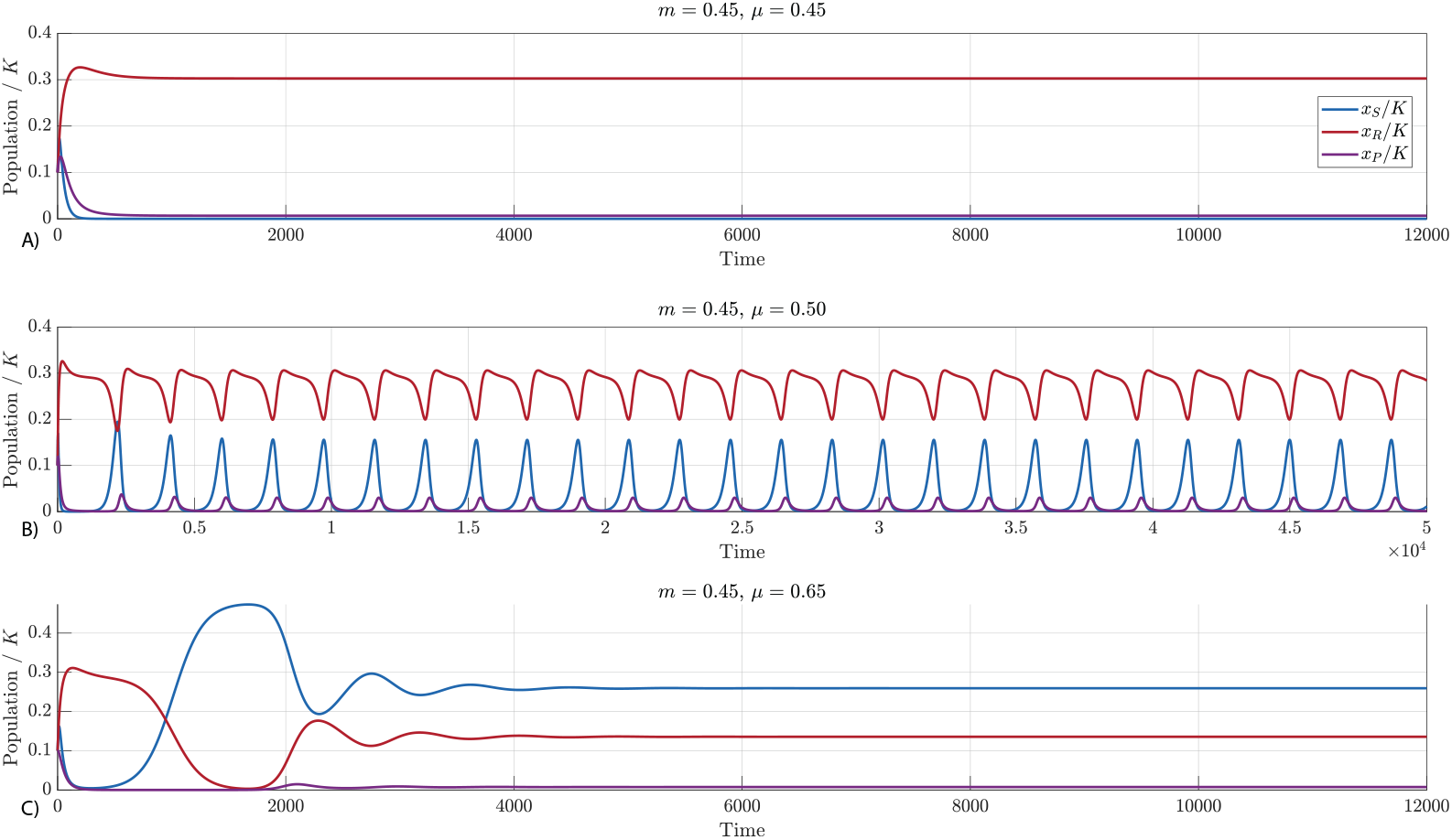
Trajectories of the original system at *m* = 0.45 with the main competition matrix and initial populations (1000, 1000, 1000). Blue, red, and purple indicate sensitive, resistant, and persister populations, respectively. Mutation values are 0.45, 0.50, and 0.65. The middle panel is integrated to time 50000; the other panels to 12000. Time is expressed in the model’s units.

For the weak-competition parameter set

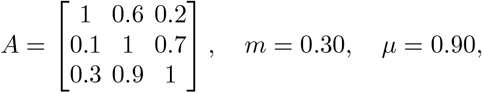

we obtain 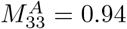. The CSR equilibrium is approximately (4164.235, 3800.350, 0) and its eigenvalues are *−*0.188, *−*0.011, and *−*0.020. The FSE and FRE are unstable, with resistant and sensitive invasion eigenvalues 0.0115 and 0.176, respectively. Thus these parameters give stable CSR, rather than two stable single-population equilibria.

Figure 7 illustrates this example and shows the CSR–CAT transcritical threshold at *µ* = 0.709.

**Fig. 7:**
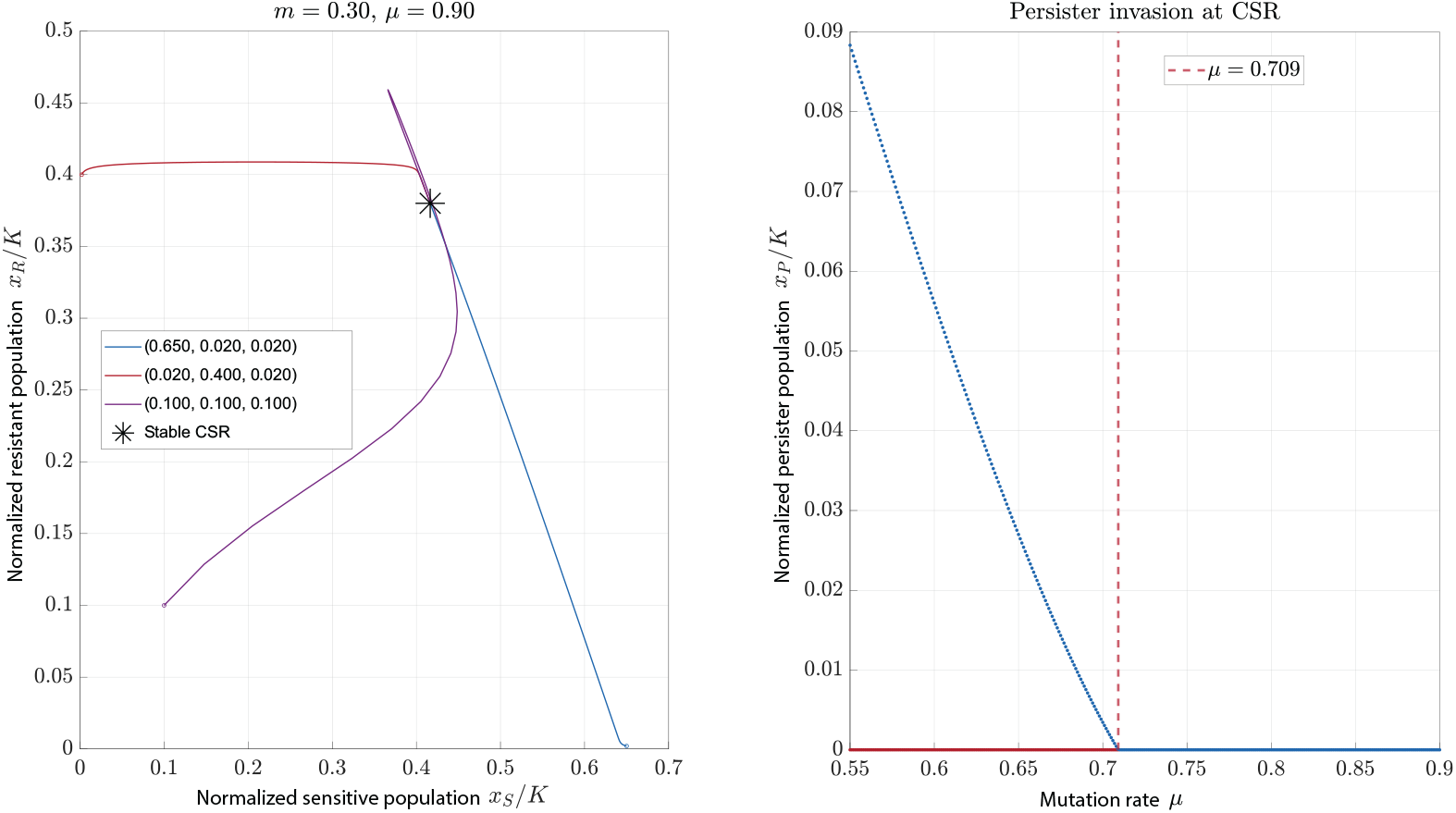
Weak-competition example. Left: trajectories from (6500, 20, 20),(20, 4000, 20), and (1000, 1000, 1000) approach the stable CSR, marked by a star. Right: the persister population on CSR and CAT branches as mutation varies, with the transcritical value 0.709 marked. Blue indicates stable and red unstable branches in the right panel. Other parameters are those in Table 2, with *u*_*R*_ = 3.3.

For the parameter values in Fig. 8, no locally stable equilibrium exists. The simulated populations continue to oscillate over the time interval shown. These observations do not establish a bound on the oscillations for all future times or demonstrate clinical safety.

**Fig. 8:**
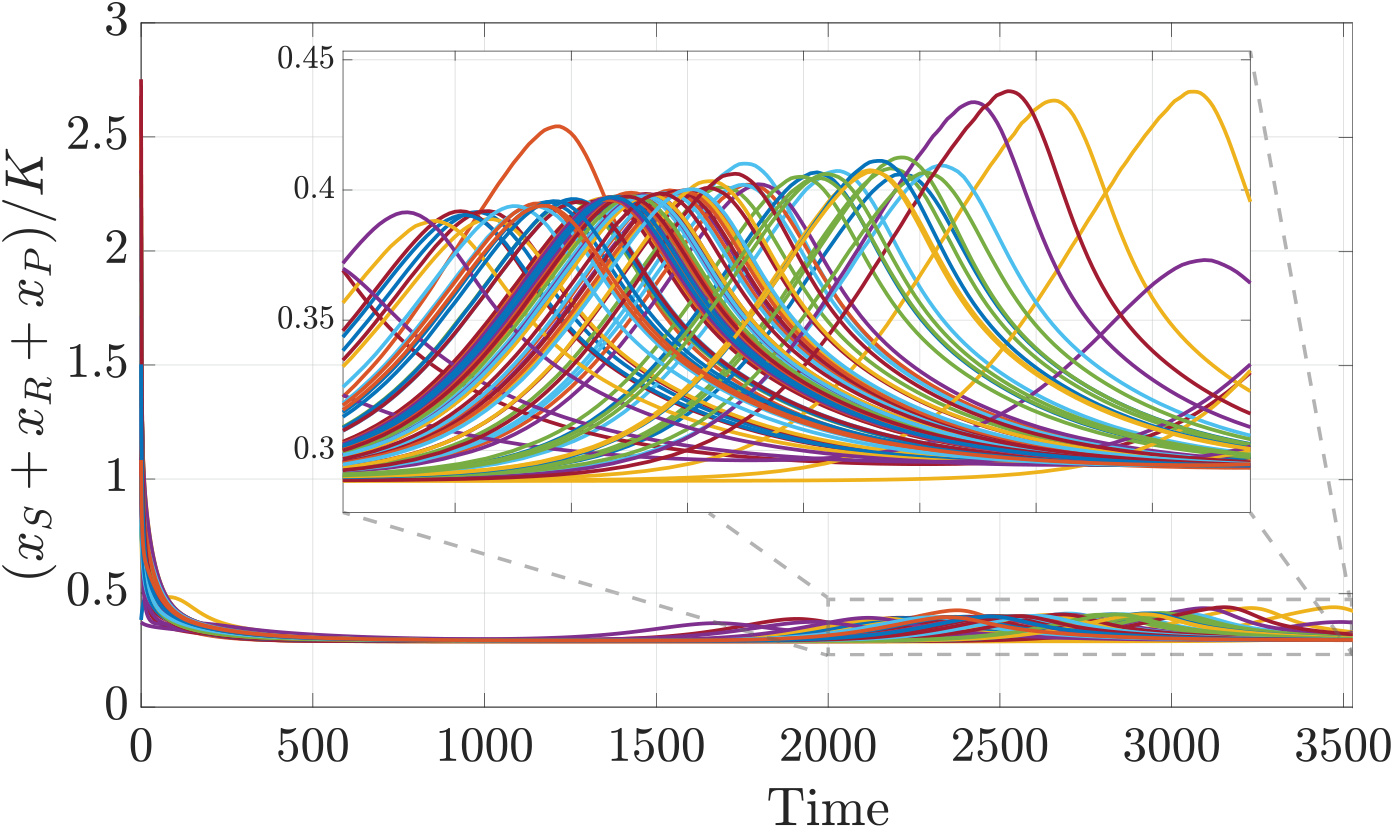
Cancer trajectories for parameter values with no locally stable equilibrium. The populations oscillate throughout the simulated time interval. The simulation does not establish a bound for all future times or clinical safety. Parameters: *m* = 0.45, *u*_*R*_ = 3.3, *µ* = 0.5, *α*_*SR*_ = 1.6, *α*_*SP*_ = 0.2, *α*_*RS*_ = 1.1, *α*_*RP*_ = 1.7, *α*_*PR*_ = 1.9, *α*_*PS*_ = 0.3.

## 5 Discussion

In this work, we developed and analyzed a mathematical model describing the evolutionary dynamics of cancer cell populations under treatment, explicitly incorporating three distinct phenotypes: sensitive, resistant, and drug-tolerant persister cells. The model extends the classical Lotka-Volterra framework by allowing irreversible transitions (mutations) from persister to resistant cells. Further, our model examines how treatment intensity, mutation rate, and inter- and intra-type competition shape the existence and stability of all possible equilibria. We derived analytical conditions under which equilibria containing one, two, or all three cell types exist and are locally stable, thereby offering theoretical insight into the long-term outcomes of stabilization therapies in heterogeneous tumors. Overall, our work extends previous research that analyzed a system of treatment-sensitive and treatment-resistant cancer cells [29].

The effect of mutation depends on treatment and competition. Stable CSR coexistence requires positive sensitive and resistant populations, *α*_*SS*_*α*_*RR*_ *> α*_*SR*_*α*_*RS*_, and a negative persister invasion eigenvalue. This product condition is also present in the two-population model [29]. Persister cells do not eliminate bistability. Simultaneous stability of the FSE and FRE requires 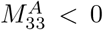 and makes a positive CSR equilibrium a saddle. The additional persister eigenvalue determines whether either single-population equilibrium remains stable in the cancer dynamic.

These findings align with and extend recent experimental and modeling studies. Single-cell lineage tracing and sequencing in EGFR-mutant lung cancer patients demonstrated that persisters can acquire resistance mutations de novo, rather than relying solely on pre-existing resistant clones [12, 24]. Further, drug exposure has been shown to elevate the mutation rate in persisters [27], enhancing their capacity to seed resistance. A recent preprint [22] builds a simulation framework around this mechanism and shows that tumors harboring even a small fraction of persisters relapse more rapidly under adaptive therapy. Similarly, bifurcation analysis by Du et al. [7] reveals that the inclusion of persister transitions can lead to oscillatory dynamics or bistable regimes, both of which complicate long-term control. These empirical and theoretical results highlight the same core conclusion as our model: that persisters undermine the assumption that resistant cells can be contained via ecological competition alone.

Our analysis shows that the stability of a persister-free equilibrium depends on whether a small persister population can grow near this equilibrium, in addition to competition between sensitive and resistant cells. The mutation thresholds and Hopf bifurcations further show that changes in persister-to-resistant mutation can alter the stable equilibrium or produce local oscillatory dynamics. These results support examining interventions that affect persister survival and the development of resistance. In this context, Sun et al. [33] identified PRMT1 as a vulnerability specific to persisters and showed that targeting it during EGFR inhibition delays resistance emergence. Corigliano et al. [4] demonstrated through optimal control that moderate, intermittent dosing strategies can suppress persister expansion more effectively than continuous therapy. These studies examine interventions that are not evaluated in our constant-treatment model. Our results provide conditions for assessing equilibrium stability under constant treatment; the efficacy of intermittent or adaptive dosing schedules requires a separate analysis with time-dependent treatment.

Our model is a simplification of biological reality. It assumes spatial homogeneity, constant mutation rates, and fixed treatment intensity. These assumptions can be relaxed in future work by incorporating spatial structure, stochastic cell fate transitions, and time-varying treatment schedules. Moreover, fitting the model to longitudinal experimental data will be essential to assess the predictive power of its equilibria and guide therapeutic design.

The bifurcation calculations identify two Hopf bifurcations and several transcritical bifurcations for the parameter sets considered. Thus, a mutation threshold can indicate invasion, a change of the attracting equilibrium, or the onset of local oscillatory dynamics. The original mutation term can become negative outside the region examined in the simulations. Therefore, the model does not preserve all nonnegative initial populations without additional restrictions. Future work should compare this formulation with a nonnegative birth-based mutation flux, estimate parameters from longitudinal measurements, and evaluate time-dependent treatment using explicit tumor-burden and toxicity criteria. Neither a stable equilibrium nor a bounded simulated oscillation alone establishes clinical safety.

Overall, our results show that persister cell survival and their transition to full resistance should be considered when assessing the existence and stability of tumor equilibria under treatment.

